# Genome assembly and isoform analysis of a highly heterozygous New Zealand fisheries species, the tarakihi (*Nemadactylus macropterus*)

**DOI:** 10.1101/2022.02.19.481167

**Authors:** Yvan Papa, Maren Wellenreuther, Mark A. Morrison, Peter A. Ritchie

**Author notes:** Corresponding author. Address: School of Biological Sciences, Victoria University of Wellington, PO Box 600, Wellington 6140, New Zealand).

## Abstract

Although being some of the most valuable and heavily exploited wild organisms, few fisheries species have been studied at the whole-genome level. This is especially the case in New Zealand, where genomics resources are urgently needed to assist fisheries management attains its sustainability goals. Here we generated 55 Gb of short Illumina reads (92× coverage) and 73 Gb of long Nanopore reads (122×) to produce the first genome assembly of the marine teleost tarakihi (*Nemadactylus macropterus*), a highly valuable fisheries species in New Zealand. An additional 300 Mb of Iso-Seq RNA reads were obtained from four tissue types of another specimen to assist in gene annotation. The final genome assembly was 568 Mb long and consisted of 1,214 scaffolds with an N50 of 3.37 Mb. The genome completeness was high, with 97.8% of complete Actinopterygii BUSCOs. Heterozygosity values estimated through k-mer counting (1.00%) and bi-allelic SNPs (0.64%) were high compared to the same values reported for other fishes. Repetitive elements covered 30.45% of the genome and 20,169 protein-coding genes were annotated. Iso-Seq analysis recovered 91,313 unique transcripts (isoforms) from 15,515 genes (mean ratio of 5.89 transcripts per gene), and the most common alternative splicing event was intron retention. This highly contiguous genome assembly along with the isoform-resolved transcriptome will provide a useful resource to assist the study of population genomics, as well as comparative eco-evolutionary studies in other teleost and related organisms.

## 1. Introduction

The tarakihi or jackass morwong (*Nemadactylus macropterus*, Centrarchiformes: Cirrhitioidei, NCBI Taxon ID: 76931) is a species of demersal marine teleost fish that is widely distributed around all inshore areas of New Zealand and along the southern coasts of Australia. It is distinguishable from other New Zealand “morwongs” by the black saddle across its nape (Roberts et al., 2015) and displays a single elongated pectoral fin ray that is characteristic of *Nemadactylus* species (Ludt et al., 2019). The species and its genus have been recently moved from the Cheilodactylidae to the Latridae following extensive revision of the taxonomy of both families, which until then was poorly understood (Kimura et al., 2018; Ludt et al., 2019). Tarakihi is an important commercial and recreational inshore fishery, especially in New Zealand, where more than 5,000 tonnes are harvested every year (Fisheries New Zealand, 2018). Like many other fisheries species, tarakihi stocks have been heavily fished over the past century. As a result, the spawning biomass is now concerningly depleted to numbers below the fisheries management soft limit of 20% on the east coast of New Zealand, where fishing effort is highest (Langley, 2018). Low effective population size and spawning biomass are of concern for the long-term sustainability of this species, particularly with added and increasing environmental pressures due to global warming. Climate change is already impacting marine ecosystems and is expected to affect the distribution and productivity of many fisheries species (Babcock et al., 2019; Burrows et al., 2011; Ramos et al., 2018).

The application of genome-wide markers for tarakihi fisheries management has been limited by the lack of a reference genome. Consequently, the first step in developing new genomic resources for this species is to assemble a high-quality reference genome that can be used to develop high-resolution markers for determining the genetic stock structure. This would offer the potential to estimate gene flow levels and detect adaptive genetic variation. Incorporating adaptive genetic variation, along with neutral variation, will greatly improve how the genetic data can be used for fisheries management (Benestan, 2019; Bernatchez et al., 2017; Papa, Oosting, et al., 2021; Thomson et al., 2021). While the neutral markers can detect reproductively isolated stocks, the adaptive loci can detect differentiation in reproductive success of migrant fish moving to locally adapted stocks. Using high-resolution markers sets for both neutral and adaptive variation has the potential to revolutionize the way genetic markers are used to define fisheries stocks.

As DNA sequencing technology is rapidly changing and improving, a range of sequencing data types has been used to produce genome assemblies, thus providing a range of genome qualities, contiguity, and completeness depending on the available technology and investment level. While short-read Illumina sequencing produces highly accurate reads, their short length (usually less than 200 bp) makes them computationally difficult to assemble. This is particularly problematic for regions that span highly repetitive segments of the genome. Complex genomes often result in highly fragmented assemblies (Koren et al., 2012; Rice & Green, 2019). The development of less accurate but long-read sequencing technologies from Oxford Nanopore and Pacific Biosciences (PacBio) has improved the assembly process by combining them with short-read data to create “hybrid”, more contiguous genome assemblies (Austin et al., 2017; Dhar et al., 2019; Jiang et al., 2019; Tan et al., 2018; Wiley & Miller, 2020; Zimin, Puiu, et al., 2017; Zimin, Stevens, et al., 2017).

The rapid improvements in sequencing technologies have also improved the ability to collect RNA sequence (RNA-seq) data. Short read RNA-seq data has been used to assist in genome annotation by first assembling a transcriptome and mapping it to the orthologous sequences to find protein-coding genes. A downside of this short read length (c. 100–150 bp) is that it is difficult or impossible to detect and characterize alternative isoforms of the coding sequences, while alternative splicing is known to occur in the vast majority of multi-exon genes (Hardwick et al., 2019). The circular consensus sequencing (CCS) PacBio technology produces reads that are both thousands of bp long and highly accurate (as opposed to the Nanopore and PacBio continuous long reads mentioned above). CCS long-read DNA sequencing can be applied to DNA (i.e. High fidelity, or HiFi reads) and RNA (i.e. isoform sequencing, or Iso-Seq). By capturing the entire sequence length of RNA molecules, Iso-Seq allows for the sequencing of complete, uninterrupted mRNAs, which enables the accurate characterization of isoforms (An et al., 2018; Byrne et al., 2019; Y. Gao et al., 2019; Hoang & Henry, 2021). Iso-Seq has been used to detect and characterize for the first time alternate splicing in the transcriptomes of several organisms, like the human (*Homo sapiens*) (Kuo et al., 2020), the chicken (*Gallus gallus*) (Kuo et al., 2017), or the goldfish (*Carassius auratus auratus*) (Gan et al., 2021). Iso-seq is now also used to annotate *de novo* genome assemblies of non-model organisms, like the cave nectar bat (*Eonycteris spelaea*) (Wen et al., 2018), the pharaoh ant (*Monomorium pharaonis*) (Q. Gao et al., 2020), the red-eared slider turtle (*Trachemys scripta elegans*) (Simison et al., 2020), or the sponge gourd (*Luffa* spp.) (Pootakham et al., 2020), allowing for the characterization of both gene functions and alternative splicing patterns.

The main goal of this study was to complete the first tarakihi genome assembly. This was achieved by using a combination of short-read Illumina and long-read Nanopore sequencing data. Four assembly pipelines were compared, three of which used algorithms implemented in MaSuRCA for hybrid assembly, and a fourth pipeline based on a trial run of low-coverage DNA sequence reads (4 Gb) generated using the PacBio HiFi platform. Iso-Seq data was used to assist with gene annotation and the identification of gene isoforms.

## 2. Materials and Methods

### 2.1 Tissue collection and nucleotide extraction

Tissues for Illumina and Nanopore sequencing were collected from a freshly vouchered *N. macropterus* specimen (standard length: 285 mm, weight: 460 g) identified as male by observation of the gonads. The specimen was a captive-bred from Plant and Food Research, Nelson, New Zealand (Figure 1A) and is thereby referred to as TARdn1 (for “tarakihi *de novo*”). A caudal fin clip and a heart piece were stored in 96% EtOH, and a kidney piece was stored in DESS (20% DMSO, 0.25 M EDTA, NaCl saturated solution). Total genomic DNA was extracted from these tissues using a high-salt extraction protocol adapted from Aljanabi & Martinez (1997) that included an RNase treatment and then suspended in Tris-EDTA buffer (10 mM Tris-HCl pH 8.0, 1 mM EDTA). The integrity of DNA fragments was assessed by gel electrophoresis in 1% agarose. The purity and quantity of DNA (concentration > 200 ng/µl, A260/280 ≈ 1.8, A60/230 ≈ 2, total weight > 20 µg) were estimated with CLARIOstar spectrometer (BMG Labtech). Purified DNA samples were sent to Annoroad Gene Technology Co. Ltd. (Beijing, China) and NextOmics Biosciences Co., Ltd. (Wuhan, China) for Illumina and Nanopore library preparation and sequencing.

**Figure 1.**
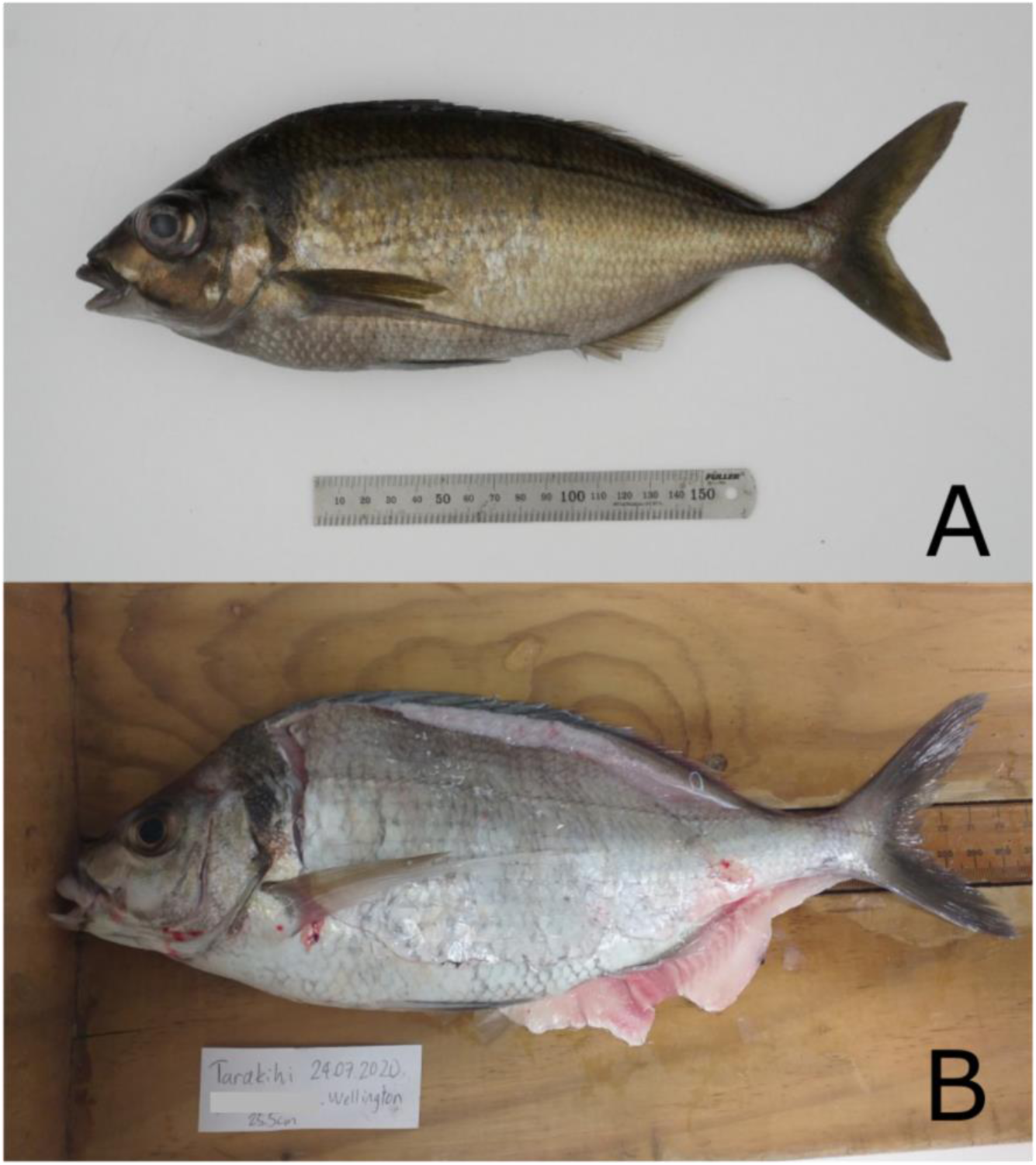
Tarakihi specimens used in this study. (A) TARdn1: captive bred specimen used for Illumina and Nanopore sequencing. (B) TARdn2: wild-caught specimen used for HiFi sequencing and Iso-Seq.

Tissues for HiFi sequencing and Iso-Seq were obtained from a wild specimen captured by a recreational fisherman at Kau Bay, in the Wellington harbour (New Zealand), thereby referred to as TARdn2 (Figure 1B). The specimen was collected for tissue sampling after being filleted by the fisherman. It had a standard length of 255 mm and was identified as male by observation of the gonads. Tissues were collected a few hours after capture and flash-frozen in liquid nitrogen. Five pieces of tissues were sent to BGI Tech Solutions Co., Ltd. (Hong Kong, China): one tissue (heart) for DNA extraction and HiFi sequencing and four tissues (liver, white muscle, brain, and spleen) for RNA extraction and Iso-Seq. DNA and RNA were extracted by BGI using a phenol-chloroform method.

### 2.2 Genome size estimation pre-sequencing

To estimate the size of the *N. macropterus* genome and ensure there was a sufficient amount of DNA sequencing for adequate coverage, genome information from closely related species was assessed. As of October 2018, only two other Centrarchiformes genome assemblies were deposited in NCBI at the scaffold level (accession numbers: GCA_002120245.1 (Murray cod, *Maccullochella peelii*), and GCA_003416845.1 (barred knifejaw, *Oplegnathus fasciatus*)), which had genome lengths of 633.24 and 766.3 Mb. Moreover, the species closest to *N. macropterus* for which genome size was estimated on the Animal Genome Size Database (http://www.genomesize.com) was the red morwong *Cheilodactylus fuscus*, with a C-value of 0.72, or approximately 700 Mb. The genome size of *N. macropterus* was thus estimated to be about 700 Mb. The quantity of Illumina and Nanopore bases to be sequenced was tuned for a deep 85× Illumina coverage (c. 60 Gb) and 140× Nanopore coverage (c. 100 Gb), following sequencing provider recommendations.

### 2.3 Library preparations and sequencing

Library preparations, sequencing, and the first filtering step (except for Nanopore reads) were performed by the sequencing providers. For Illumina reads, DNA samples were sheared with Bioruptor® Pico system (Diagenode) for a fragment insert size of 350+/-50 bp, and a PCR-free library was obtained with NEBNext® Ultra™ II DNA Library Prep Kit for Illumina (New England Biolabs). Approximately 200 million of 150 bases pair-end reads were generated using the HiSeq X System (Illumina). Raw Illumina reads were filtered by discarding read pairs if (1) one read contained some adapter contamination for more than five nucleotides, (2) more than 10% of bases were uncertain in either one read, or (3) the proportion of bases with Quality Value ≤ 19 was over 50% in either one read. For Nanopore library preparation, large size DNA fragments were selected by automated gel electrophoresis with BluePippin (Sage Science) followed by enrichment and purification using beads. Fragmented DNA was then end-repaired, A-tailed, and purified, and adapter ligation was done using the Ligation Sequencing Kit 1D 108 (Oxford Nanopore Technologies). The resulting DNA library of 20–40 Kb fragments was then loaded into two flow cells for real-time single-molecule sequencing on PromethION (Oxford Nanopore Technologies). Reads were base-called from their raw FAST5 files using Albacore 2.0.1 (https://community.nanoporetech.com). HiFi library was prepped with SMRTbell® Express Template Prep Kit 2.0 (Pacific Biosciences) and CCS was performed on one-third of an SMRT Cell 8M with a PacBio Sequel II sequencer. ZMWs were filtered to retain a minimum of three passes and a predicted quality value (RQ) of 99. Four Iso-Seq libraries of 0– 5 kb insert sizes (one per tissue) were generated using the SMRTbell® Express Template Prep Kit 2.0. The multiplexed libraries were sequenced on one SMRT Cell 8M with a PacBio Sequel II sequencer, resulting in 3.6 million polymerase reads from which sub-reads were extracted.

### 2.4 Illumina reads: Quality and contamination filtering

Primary quality filtering resulted in 405.2 million Illumina pair-end reads (60.78 Gb). Quality metrics of these filtered reads were visualized with FastQC v0.11.7 (Andrews, 2018) before proceeding to the next steps. Kraken v2.0.7-beta (Wood et al., 2019) was used to detect and filter contamination from archaea, bacteria, viral, and human sequences based on the MiniKraken2 v2 8GB database (Wood, 2019). The 9.25% of reads that were classified as contaminants were discarded, leading to 367.8 million non-contaminated reads (55.16 Gb) (Table 1).

**Table 1.** Summary of number, base quantity, and length of reads obtained at several steps of the quality filtering pipelines.

| Reads | Number of reads | Total number of bases | Minimum read length | Average read length | Maximum read length |
| --- | --- | --- | --- | --- | --- |
| Raw Illumina PE reads | 425,740,632 | 63,861,094,800 | 150 | 150 | 150 |
| Quality-filtered Illumina PE reads | 405,228,300 | 60,784,245,000 | 150 | 150 | 150 |
| Uncontaminated Illumina PE reads | 367,760,592 | 55,164,088,800 | 150 | 150 | 150 |
| **Final Illumina PE reads** | 366,065,036 | 54,909,755,400 | 150 | 150 | 150 |
| Raw Nanopore reads cell 1 | 8,270,853 | 52,169,467,195 | 5 | 6,307.6 | 1,029,695 |
| Raw Nanopore reads cell 2 | 7,229,556 | 47,015,342,634 | 5 | 6,503.2 | 1,035,919 |
| **Final Nanopore reads** | 9,178,726 | 73,394,980,774 | 450 | 7,996.2 | 182,445 |
| HiFi reads | 285,997 | 4,009,988,664 | 49 | 14,021.1 | 27,427 |
| Iso-Seq sub-reads (4 tissues) | 171,924,197 | 302,196,904,697 | 51 | 2,601.15 | 278,803 |
| **Final Iso-Seq transcripts** | 91,602 | 312,308,038 | 80 | 3,409.4 | 10,426 |

Notes: Reads in bold were the ones used in the final retained (Flye polished) assembly. Final Illumina PE reads have been filtered for quality, DNA contamination, and mitochondrial DNA. Final Nanopore reads have been filtered for quality. Final Iso-Seq CCS transcripts were filtered for quality and repeat transcripts and were non-redundant.

### 2.5 Illumina reads: Mitogenome assembly and exclusion

Illumina reads filtered for quality and contamination were mapped against the Peruvian morwong (*Cheilodactylus variegatus*) complete mitochondrial sequence retrieved from Genbank (accession number: KP704218.1) with Geneious v11.04 (Kearse et al., 2012) using five iterations of the default mapper set to the highest sensitivity. The extracted consensus sequence resulted in a 16,650 bp assembly of the *N. macropterus* mitogenome. The mitogenome was then annotated using the MitoAnnotator web interface (Iwasaki et al., 2013). Sequences of mitochondrial origin were then filtered out of the Illumina reads as follows: first, bwa-kit v0.7.15 (Li & Durbin, 2009) was used to align the Illumina reads to the indexed *N. macropterus* reference mitogenome with default parameters. Among other aligners, the BWA-MEM algorithm (Li, 2013) was selected because it is the most accurate for this type of short-read data (Keel & Snelling, 2018). In the resulting SAM alignment, 0.46% of reads mapped to the mitogenome. Then, all the reads from the alignment that did not map to the mitogenome were extracted to a new mitochondria-free alignment using SAMtools v1.9 (Li et al., 2009) `view` with parameters `-b -f 4`, sorted by name, and finally converted back to FASTQ paired-end reads with bedtools v2.27.1 (Quinlan & Hall, 2010).

### 2.6 Genome size estimation post-sequencing

Genome size and sequencing coverage based on the Illumina sequence reads was performed with a *k*-mer frequency analysis. Total number of 17, 21, and 27-mers were counted with jellyfish v2.2.10 (Marçais & Kingsford, 2011) command `count` and the resulting histograms were computed with command `histo`. The histograms were analyzed with GenomeScope (Vurture et al., 2017), which estimated a genome size of c. 516–520 Mb, with a high heterozygosity level of 1.01–1.07 % and a duplication level of 0.98–1.10 % (Figure 2, Supplementary Figure 1). This estimated haploid genome size was consistent, albeit c. 150 Mb lower than the size estimated pre-sequencing. However, it is common for *k*-mer estimated size and genome assembly size to be smaller than the size estimated with C-value (Austin et al., 2017; Feron et al., 2020; Jansen et al., 2017). The heterozygous coverage of 40× was considered sufficient for performing genome assembly.

**Figure 2.**
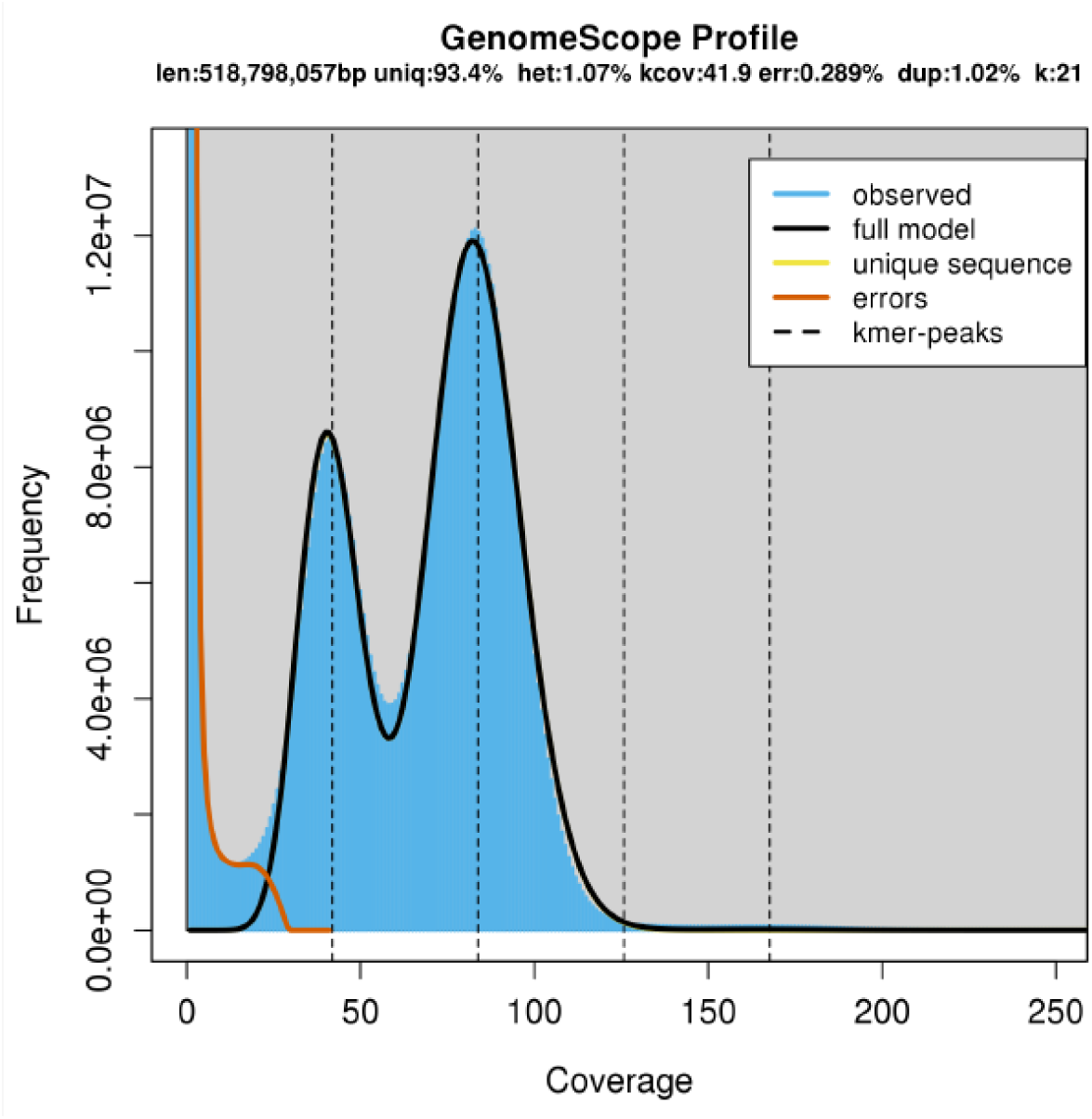
Histogram of 21-mer frequency in Illumina reads. Estimation of genome size, heterozygosity, and duplicated regions. The first and second peaks show the *k*-mer frequency of heterozygous and homologous regions, respectively. See Supplementary Figure 1 for 17- and 21-mer models.

### 2.7 Nanopore reads sequencing and filtering

A total of 99.18 Gb was obtained from the raw unfiltered reads, with an average read length above 6 Kb and a maximum length above 1 Mb (Table 1). Quality control of the raw reads was performed with NanoPack v1.0.0 (De Coster et al., 2018) using `NanoStat` on both FASTQ reads and Albacore summary files. Nanopore reads were filtered and trimmed with `NanoFilt` by applying a minimum length cut-off of 500 bases (Tan et al., 2018), a minimum average read quality score of 7 (c. 80% base call accuracy), and removing the first 50 nucleotides following the author’s recommendations (De Coster, 2017). Given that quality values based on summary files were slightly lower overall than when based on reads (as expected, c.f. github.com/wdecoster/nanofilt), the quality filtering was done based on the summary file values to be more stringent. Filter-trimmed reads from both cells were merged into a single FASTQ file.

### 2.8 Illumina + Nanopore hybrid assembly

*De novo* genome assembly of short and long reads was performed with the Maryland Super-Read Celera Assembler pipeline, MaSuRCA (Zimin et al., 2013; Zimin, Puiu, et al., 2017). This is one of the most common assemblers for performing short and long reads hybrid genome assemblies of eukaryotes, with consistently good results across studies (Jiang et al., 2019; Tan et al., 2018; Thai et al., 2019). In brief, MaSuRCA typically works as follows: Illumina paired-end short reads are first assembled into non-ambiguous super-reads, which are then mapped to Nanopore reads to further assemble them in long, high-quality pre-mega-reads. If there are gaps between mega-reads in respect to their mapping to the Nanopore reads, these gaps are filled with the Nanopore read sequence only if the Nanopore read stretch meets some minimum criteria of coverage and quality to produce the mega-reads. If there are still gaps that cannot be merged between mega-reads due to poor quality of the Nanopore sequence, regions flanking these gaps are linked together as linking pair mates. The mega-reads and linking pairs are then assembled with either CABOG or Flye (see below).

Before assembly, the filtered Illumina reads were not trimmed or edited as per MaSuRCA author recommendation (https://github.com/alekseyzimin/masurca). The hybrid Illumina + Nanopore assembly was run on MaSuRCA v3.2.9 with recommended parameters, automatic *k*-mer size computation, and a jellyfish hash size of 20,000,000,000 (PE = pe 350 50, NANOPORE, EXTEND_JUMP_READS = 0, GRAPH_KMER_SIZE = auto, USE_LINKING_MATES = 0, USE_GRID = 0, GRID_BATCH_SIZE = 300000000, LHE_COVERAGE=25, MEGA_READS_ONE_PASS=0, LIMIT_JUMP_COVERAGE = 300, CA_PARAMETERS = cgwErrorRate = 0.15, KMER_COUNT_THRESHOLD = 1, CLOSE_GAPS = 1, NUM_THREADS = 32, JF_SIZE = 20000000000, SOAP_ASSEMBLY = 0). MaSuRCA v3.2.9 uses a modified version of the CABOG assembler (Miller et al., 2008) for the final assembly of corrected mega-reads. However, later releases of MaSuRCA included the Flye assembler (Kolmogorov et al., 2019) as a supposedly faster and more accurate alternative tool for the same step. To compare both methods, a second assembly was run on MaSuRCA v3.4.1 with the same parameters as above, but this time using FLYE_ASSEMBLY = 1. The Flye assembly was subsequently polished with POLCA (Zimin & Salzberg, 2020) as implemented in MaSuRCA v3.4.1 on default settings, using the clean Illumina reads to fix substitutions and indel errors.

### 2.9 HiFi sequencing and assembly

HiFi reads were converted from BAM to FASTA and FASTQ with SMRTLink v9.0 (PacBio, 2020) `bam2fastx`. Assembly was performed with hifiasm v0.13 (Cheng et al., 2021) using default parameters. The primary contigs were extracted from the GFA graph and converted to FASTA with command `awk ’/^S/{print “>”$2;print $3}’`. Another assembly was also tentatively performed with HiCanu as implemented in Canu v2.1.1 (Nurk et al., 2020), with an estimated genome size of 600 Mb. However, the read coverage estimated (6.68×) was lower than the minimum coverage allowed by HiCanu (10×), so the assembly could not be completed.

### 2.10 Quality assessment and comparison of assemblies

After each assembly, basic contiguity statistics were computed with bbmap v38.31 (Bushnell, 2018) script `stats.sh`. Length, GC content, and GC skew of scaffolds in all assemblies were also reported with seqkit v0.10.1 (Shen et al., 2016) command `fx2tab`. To assess the completeness of the assemblies, the Benchmarking Universal Single-Copy Orthologs (BUSCO) tool v3.0.2 (Simão et al., 2015) was used with parameter `-sp zebrafish` on the Actinopterygii odb9 orthologs set, which contains 4,584 single-copy orthologs that are present in at least 90% of ray-finned fish species. Augustus v3.3.1 (Stanke et al., 2004), NCBI blast+ v2.7.1 (Camacho et al., 2009), hmmer v3.2.1 (Eddy, 2011), and R v3.6.0 (R Core Team, 2020) were also required to run the BUSCO shell script.

The quality of the CABOG and Flye assemblies was further compared by mapping clean Illumina reads back to the assemblies themselves with bwa-kit v0.7.15 using `bwa mem -a-M`. The resulting alignment files were also used to plot Feature Response Curves (FRC) (Vezzi et al., 2012b) with FRCbam v5b3f53e-0 (Vezzi et al., 2012a). This allowed comparison of quality of the assemblies without relying on contiguity, by plotting the accumulation of error “features” along the genome (e.g. areas with low or high coverage, number of unpaired reads, misoriented reads). The presence of unmerged haplotigs in the CABOG and the Flye polished assembly was investigated by using minimap v2.16 (Li, 2018) with parameters `-ax map- ont --secondary = no` to map the clean Nanopore reads back to the assembly and then analyzing the resulting alignment with Purge Haplotigs v1.1.1 (Roach et al., 2018) command `hist`. The presence of trailing Ns in the Flye polished assembly was tested by using seqkit v0.10.1 command `-is replace -p “^n+|n+$” -r “”` and comparing the input and the output.

A last quality check of the CABOG and Flye polished assemblies was done by plotting assemblies against each other and against two chromosome-level fish assemblies using MashMap 2.0 (Jain et al., 2018) with a minimum mapping segment length of 500 bp and a minimum identity of 85% (for comparison between tarakihi assemblies) and 90% (for comparison between different species). To visualize the presence of potential misassemblies on the longest scaffolds, the results from MashMap were used to plot the mappings of these scaffolds between different assemblies with a custom R script (plot_mashmap_scaffolds.R). The first fish chromosome-level assembly used for comparison was the mandarin fish *Siniperca chuatsi* (SinChu7, GCA_011952085.1) because it was the phylogenetically closest chromosome-level assembly (Centrarchiformes, Centrarchoidei) available on NCBI at the time this analysis was performed. The second was the Australasian snapper *Chrysophrys auratus* (SNA1, https://www.genomics-aotearoa.org.nz/data), in order to compare with a well-curated specimen from a more evolutionarily distant species.

Final visualization of contiguity and completeness of the genome assemblies was generated with assembly-stats v17.02 (Challis, 2017) as implemented in the grpiccoli container (Piccoli, 2021).

### 2.11 Estimation of heterozygosity

The heterozygosity of TARdn1 was estimated a second time by calling SNPs from the Illumina reads aligned to the final assembly. The reads were mapped to the polished assembly with bwa-kit v0.7.15 using the command `bwa mem -a -M`. Duplicates were marked with picard v2.18.20 (Broad Institute, 2019) `MarkDuplicates`. SNPs were called using bcftools v1.9 (Li, 2011) commands `mpileup` (`–C50 –q10 –incl-flags 2`) and `call` (`-m -- variants-only -- skip-variants indels`). To filter for good quality SNPs, variants depth distribution was plotted. The modal depth of coverage was 82, with an increase in steepness starting at c. 20 and a decrease starting at c. 120 (Supplementary Figure 2). Consequently, the final SNP set was filtered with vcftools v0.1.16 (Danecek et al., 2011) for a minimum reference allele frequency of 0.25, a genotype depth of minimum 20 and maximum 120, and a minimum site quality of 20.

### 2.12 Genome repetitive elements detection

Repetitive elements (RE) in the *N. macropterus* genome were identified both by *de novo* modeling and based on repeats homology. RepeatModeler v2.0.1 (Flynn et al., 2020), as implemented in Dfam TE Tools container v1.2 (https://github.com/Dfam-consortium/TETools), was used to identify repeat models *de novo* using parameter `- LTRStruct` to include the detection of long terminal repeat retrotransposons. For the homology-based library, RepeatMasker v4.1.1 (Smit et al., 2013) tool `famdb.py` was used to obtain known Actinopterygii repeats from the combined total Dfam v3.3 (Storer et al., 2021) and RepBase RepeatMasker Edition v20181026 (Bao et al., 2015) databases, using parameters `--ancestors –descendants --include-class-in-name --add- reverse-complement`. Both *de novo* and homology-based repeat libraries were then concatenated in a custom repeat library for *N. macropterus*. The genome assembly sequences were then mapped against the custom repeat library with RepeatMasker v4.1.1 (`-gff - xsmall`) to classify repeat regions, create a repeat annotation file, and produce a “soft-masked” (i.e. masked bases in lower case) genome assembly. An alternate “hard-masked” assembly was also created by converting lower cases in the soft-masked assembly into Ns.

### 2.13 Iso-Seq analysis

Iso-Seq sub-reads were processed with the SMRTLink v9.0 Iso-Seq pipeline. Circular consensus sequences were generated from the sub-reads with command `ccs` using a minimum read quality (RQ) of 0.9. Clontech and NEB primers removal and de-multiplexing were performed using `lima` with parameters `--isoseq --dump-clips --peek- guess`. Poly-A tails were trimmed and concatemers were removed with `isoseq3 refine`. At that point, BAM files containing sequence reads from the four tissues were merged in one. Clustering and polishing of full-length reads were performed with `isoseq3 cluster` and parameter `--use-qvs` to obtain a dataset of high-quality isoforms with a predicted accuracy > 0.99. These high-quality polished isoforms were then aligned to the unmasked *N. macropterus* genome with pbmm2 (`--preset ISOSEQ --sort`).

Subsequently, redundant isoforms were collapsed into non-redundant transcripts loci using the command `collapse`. Non-redundant transcripts were screened for REs against the *N. macropterus* custom repeat library with RepeatMasker v4.1.1. Transcripts with ≥ 70% bases masked were considered REs. Identified REs were discarded from further analyses using a custom bash script for filtering (Count_filter_N_isoseqrepeats.bash) and categorized using a custom R script (R_charachterize_transcripts.R).

Alternative splicing (AS) events in the repeat-cleaned Iso-Seq reads were counted and classified with SUPPA v2.3 (Trincado et al., 2018) with default parameters. These results were compared with reported AS values for other animal species from studies that also used SUPPA on Iso-Seq reads. Results reported were compiled for the zebrafish (*Danio rerio*) (Nudelman et al., 2018), the goldfish (*Carassius auratus auratus*) (Gan et al., 2021), the Wuchang bream (*Megalobrama amblycephala*) (Chen et al., 2021), the whiteleg shrimp (*Litopenaeus vannamei*) (X. Zhang et al., 2019), and the cave nectar bat (*Eonycteris spelaea*) (Wen et al., 2018).

### 2.14 Genome annotation

The unmasked *N. macropterus* genome was annotated using the MAKER v2.31.10 (Holt & Yandell, 2011) pipeline. First, the simple repeats were filtered out of the repeats annotation file with a custom bash script (rm_simple_repeats.bash) to retain only complex repeats. Only complex repeats were kept because MAKER will hard-mask every region provided in the repeats annotation file before running, discarding them from the gene detection process. However, simple repeats should be available for gene annotation because low-complexity regions are expected within many genes. Hard-masking only complex repeats regions as a first step allows MAKER to subsequently identify and soft-mask the simple repeats regions internally. Gene matches that start in a non-masked region but extend in a soft-masked region can then be taken into account in the gene detection process. A first round of MAKER was run on the unmasked genome using the high-quality, non-redundant, non-repetitive Iso-Seq transcripts to infer gene predictions (`est2genome = 1`). For repeat masking during this step, the complex repeats GFF file was provided for hard masking and only simple repeats were annotated (`model_org = simple`). All GFF and FASTA outputs were then merged with `ggf3_merge` and `fasta_merge`. Training files for the *ab initio* gene predictors SNAP v2013.11.29 (Korf, 2004) and Augustus v3.3.1 (Stanke et al., 2004) were generated based on round 1 results. For SNAP, only gene models with a maximum Annotation Edit Distance (AED) of 0.25 and a minimum protein length of 50 were used. For Augustus, all the regions that contain mRNA annotations, including the 1,000 surrounding bp, were extracted to a FASTA file using a custom bash script (augustus_rndx.bash). BUSCO v3.0.2 was then run in “genome” mode on the FASTA file using the Actinopterygii odb9 orthologs set, the zebrafish as initial HMM model, and parameter `--long` to self-train Augustus. MAKER was then run a second time using SNAP and Augustus training files, as well as the Iso-Seq transcriptome and repeats alignments as evidence (`est2genome = 0`). For this, all lines containing “est2genome” and “repeat” in the merged GFF from round 1 were extracted and copied in two files that were provided as evidence with the parameters `est_gff` and `rm_gff`, respectively. Additionally, gene predictions were also inferred from protein homology during this round (`protein2genome = 1`), by using protein sequences of zebrafish (*Danio rerio*), three-spined stickleback (*Gasterosteus aculeatus*), spotted gar (*Lepisosteus oculatus*), Nile tilapia (*Oreochromis niloticus*), medaka (*Oryzias latipes*), Japanese puffer (*Takifugu rubripes*), green spotted puffer (*Tetraodon nigroviridis*), and southern platyfish (*Xiphophorus maculatus*) that were downloaded from Ensembl release version 103 (Kersey et al., 2016). After that, SNAP was trained again using the results from round 2, and a third run was performed by using the *ab initio* training files, as well as the extracted repeats, Iso-Seq, and protein homology GFF files as evidence. Genes were renamed with MAKER `maker_map_ids` and `map_x_ids`.

All proteins predicted from the second round of MAKER were blasted against the NCBI non-redundant protein sequences database (NR) with blastp (`-evalue 1e-6 -max_hsps 1-max_target_seqs 1 -outfmt 6`) as implemented in blast+ v2.6.0. All putative gene functions based on the best homology matches were annotated in the genome with a custom bash script (add_blast_annotation_custom.bash). Protein-coding genes were also searched for protein domains and signatures and annotated for InterPro (IPR), Pfam, and Gene Ontology (GO) terms using InterProScan v5.50-84.0 (Jones et al., 2014) and MAKER `ipr_update_gff`. Protein domains were exported as features in a GFF file using MAKER `iprscan2gff3`.

Finally, low-quality genes were identified with AGAT v0.6.0 (Dainat, 2021). These genes were filtered out if they were shorter than 50 amino acids and flagged if they had an incomplete open reading frame (ORF). Gene models produced by the second MAKER round were kept as the final reference dataset based on their higher number, AED distribution, and BUSCO completeness (Supplementary Table 2). Genome annotation was also inspected visually with JBrowse v1.1.10 (Skinner et al., 2009).

### 2.15 General bioinformatics tools

After each assembly, scaffolds were sorted by size using seqkit v0.10.1 command `sort -l -r -2` and renamed with command `replace -p .+ -r “{nr}”` (i.e. scaffold “1” being the longest, etc.). All alignment files were systematically sorted by leftmost coordinates, converted to BAM, and indexed with SAMtools v1.9. Alignment summary reports were produced with BAMtools v2.5.1 (Barnett et al., 2011). FASTQ files were converted in FASTA when needed with seqtk v1.3 (https://github.com/lh3/seqtk), and similarly, GFFs were converted to GTF with AGAT v0.6.0. Analyses were performed on Rāpoi, the Victoria University of Wellington high-performance computer cluster. Analyses requiring R scripts were performed in R v4.02 (R Core Team, 2020) on RStudio (RStudio Team, 2020). All bash and R scripts used for this chapter are available on GitHub on the following repository: https://github.com/yvanpapa/tarakihi_genome_assembly.

## 3. Results

### 3.1 Genome sequencing

Illumina sequencing reads filtering (i.e. quality, contamination, and mitochondria) resulted in a final dataset of 54.91 Gb short reads (Table 1) with a c. 92× depth of coverage. The GC content was 43% and the overall sequence read quality was high. Both forward and reverse reads passed all the FastQC criteria, i.e. they were never flagged for poor quality (Supplementary Figure 3). Although there was a small bias in per base sequence contents of the first c. 10 bases, this was expected due to the non-random nature of the hexamer priming step during sequencing (Hansen et al., 2010). This slight deviation from uniformity in sequence content was not considered an issue because there is no quantitative step involved in the analyses based on the short reads. Nanopore sequencing, filtering, and trimming resulted in 9.18 million reads (73.39 Gb), or 122× coverage, with an average read length of 8 Kb (Table 1), a mean read quality of 7.9, and an N50 length of 9.5 Kb. A total of 285,997 CCS Hi-Fi reads (4.01 Gb) and 91,602 repeat-free, non-redundant, high-quality Iso-Seq transcripts (312.31 Mb) were also obtained.

### 3.2 Assemblies comparison and quality assessment

The Flye assembly reduced the number of scaffolds by more than half compared to the CABOG assembly (Table 2). The scaffold N50 length of the Flye assembly was almost twice as long and the number of complete BUSCOs was higher. The Flye assembly size was also more consistent with the haploid genome size pre-estimated by *k*-mer counting (c. 520 Mb) than the CABOG assembly. Interestingly, the Flye assembly also corrected a misassembly of the first scaffold of the CABOG assembly (see below). Polishing the Flye assembly resulted in the correction of 43,080 substitutions errors and 42,783 deletion errors. The polished assembly had the same number of scaffold and contigs, but a few hundred fewer bases, and one missing BUSCO was recovered into an additional single-copy BUSCO. The hifiasm assembly performed on the HiFi reads did not produce satisfactory results compared to the Illumina + Nanopore hybrid assemblies, with six to ten times more scaffolds, an N50 length 50 times smaller, and a BUSCO completeness lower than 90%. This was most probably due to the low coverage of HiFi reads (c. 6.5x) used for this sequencing trial.

**Table 2.**
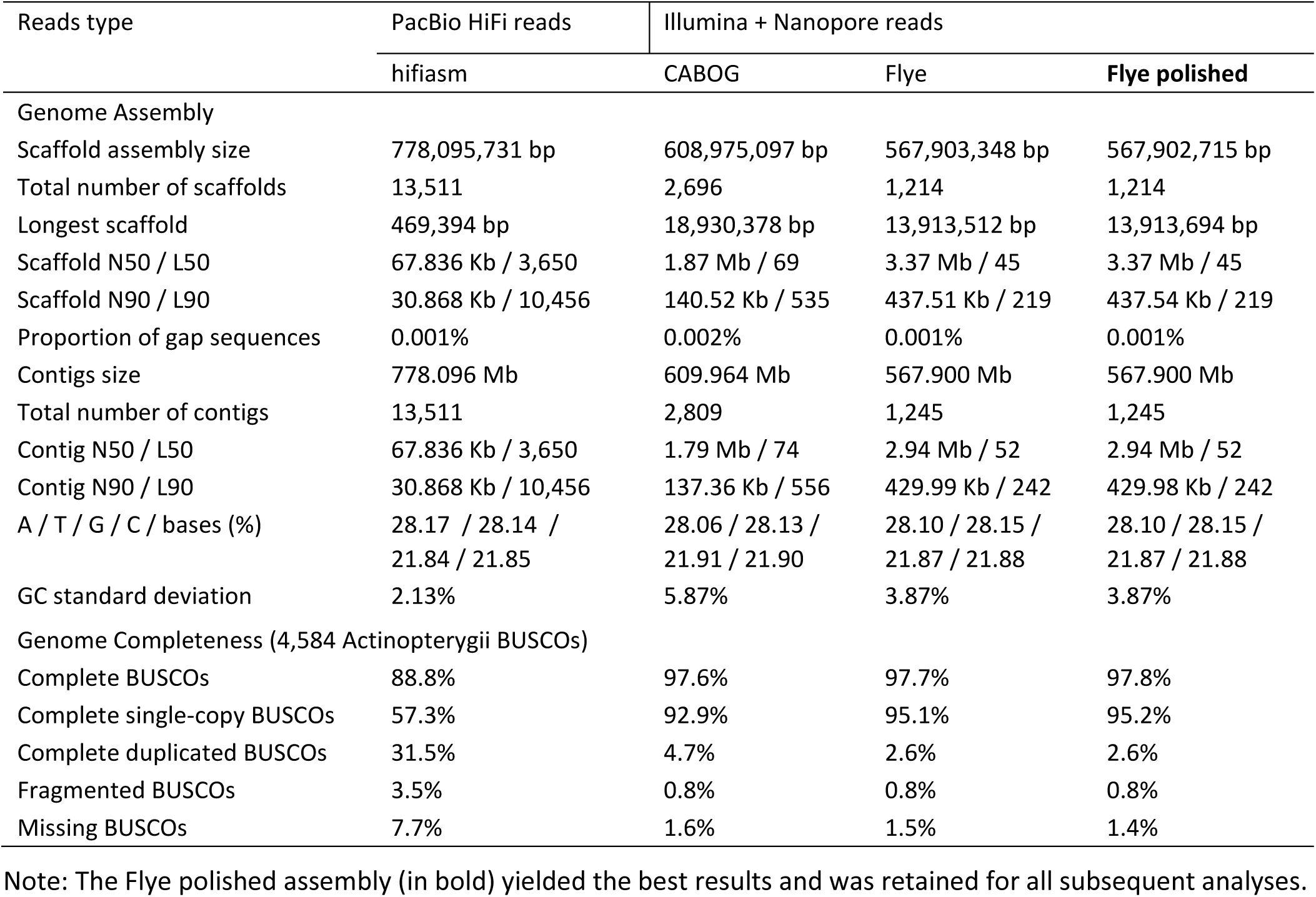
General statistics of the four assemblies produced.

Approximately 99.7% of Illumina reads could be mapped back to the CABOG assembly, and 99.8% to both Flye assemblies, making the Flye assemblies slightly more accurate according to that metric. The Flye polished assembly had a slightly higher proportion of “proper-pairs” reads mapped (86.23%) than the un-polished assembly (85.7%). FRC curves showed that both Flye assemblies were more accurate than the CABOG assembly (Figure 3). Moreover, while both the unpolished and polished Flye assemblies have a very similar curve, for the same genome coverage, the polished Flye assembly always had a slightly lower amount of cumulative errors compared to the un-polished assembly (Supplementary Figure 4.).

**Figure 3.**
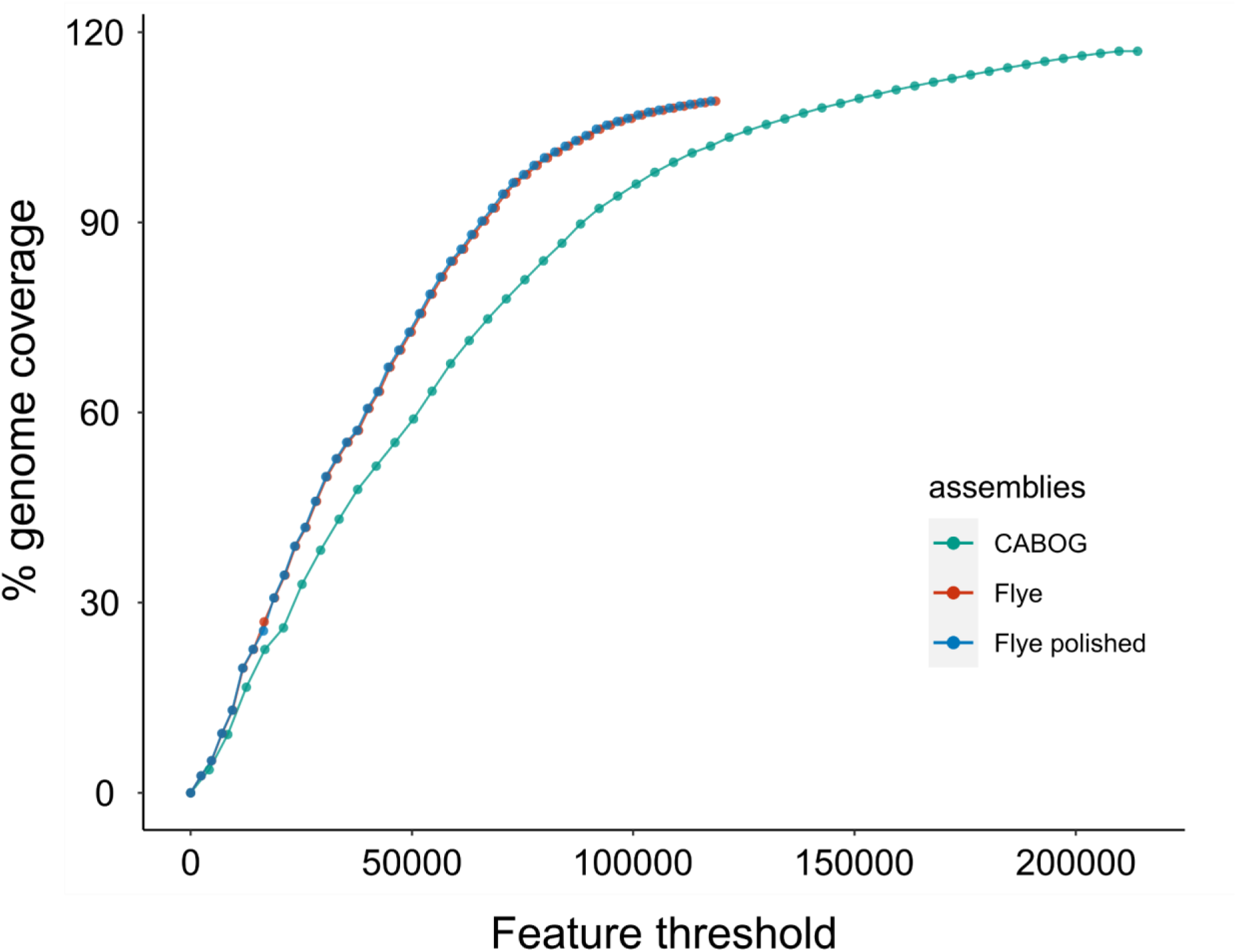
FRC curves for the CABOG, Flye, and Flye polished assembly. The Y-axis represents the cumulative size of the assembly and the X-axis is the cumulative number of potential errors (i.e. “features”). Assemblies for which the curves are steeper are considered more accurate.

While there was evidence of the presence of unmerged haplotigs in the CABOG assembly (Figure 4A), none were detected in the Flye polished assembly (Figure 4B), thus a filtering step was not required. Trailing Ns were not present in the Flye polished assembly either.

**Figure 4.**
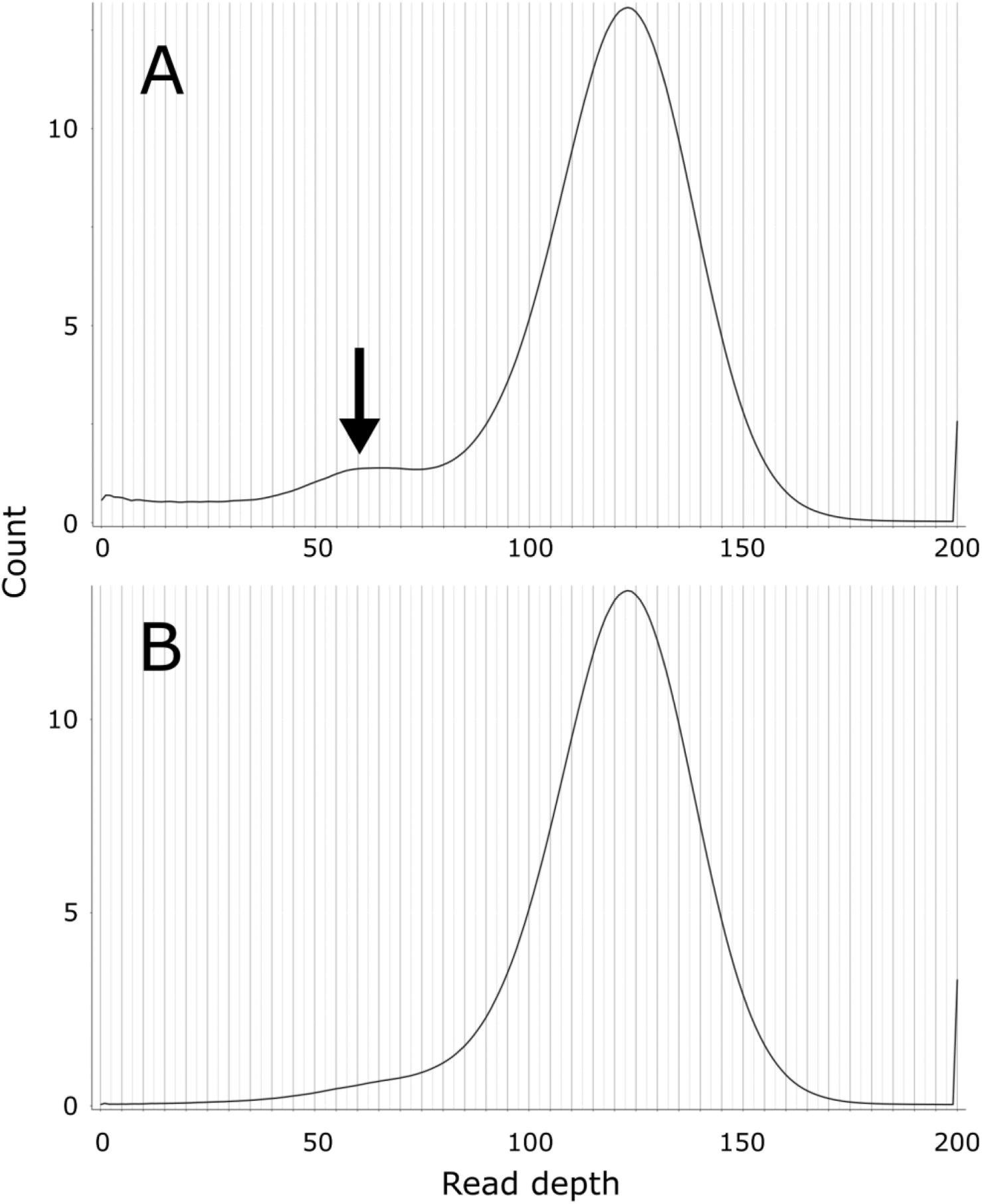
Read depth histograms of the genome assemblies contigs, obtained by mapping the clean Nanopore reads back to the assembly. A unimodal distribution with a peak equal to the sequencing reads depth is expected for a haplotig-free assembly. Another peak at half of the sequencing reads depth (arrow) is indicative of the presence of unmerged haplotigs. A: CABOG assembly B: Flye polished assembly.

Interestingly, the longest scaffold of the CABOG assembly, scaffold 1, was 5 Mb longer than the longest scaffold of the Flye assembly (Table 2). Between-scaffolds alignment scores obtained from MashMap (Supplementary Figure 5) were used to visualize a potential misassembly at that scaffold. The longest scaffold of the CABOG assembly corresponded indeed to the two longest scaffolds of the polished Flye assembly, scaffolds 1 and 2 (Figure 5A). The CABOG scaffold 1 is highly likely to have been misassembled since it also corresponds to two long regions in two different linkage groups (i.e. chromosomes) in both chromosome-level assemblies of *S. chuatsi* and *C. auratus.* This is not the case for scaffold 1 in the polished Flye assembly (Figure 5B). This supported the interpretation that the “correct” longest scaffold is the one from the polished Flye assembly.

**Figure 5.**
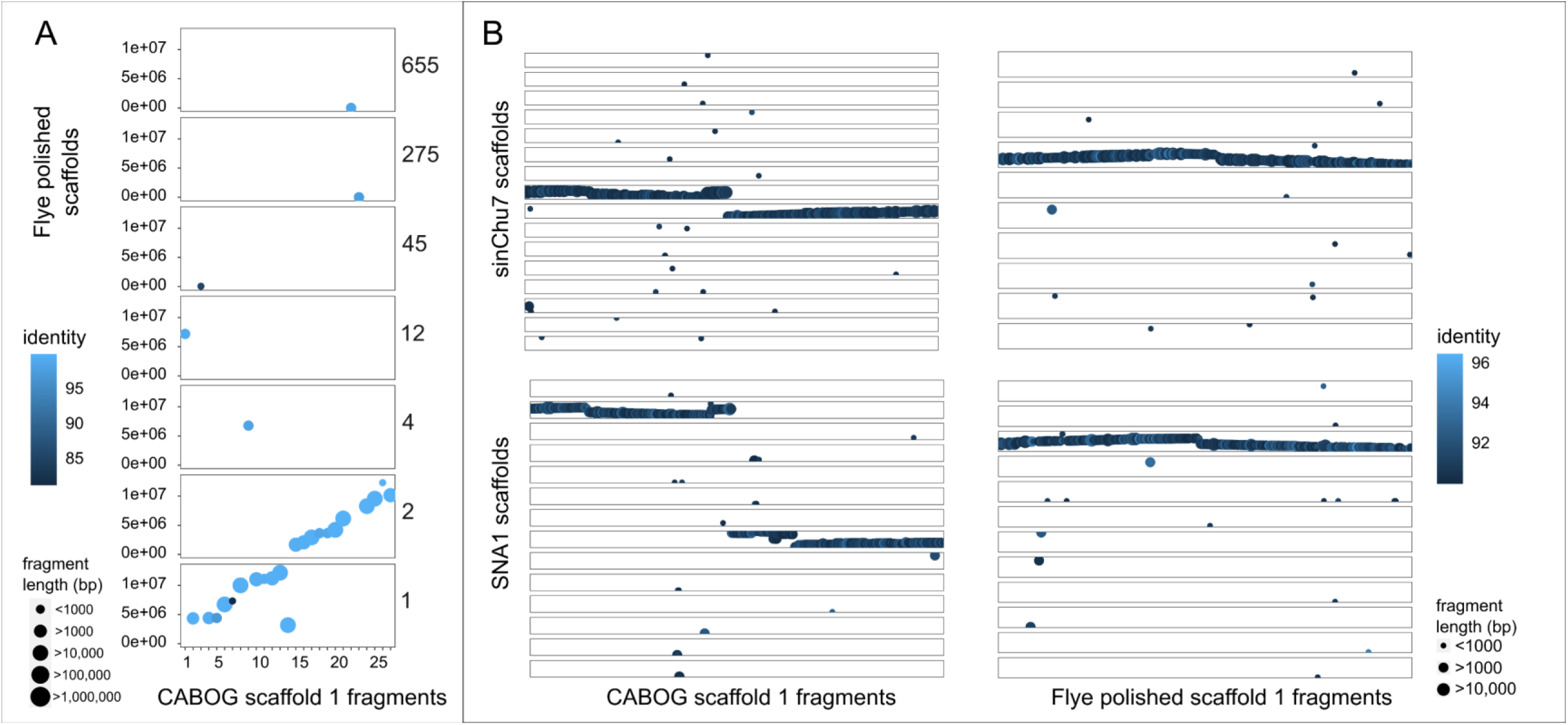
Scaffolds plotted against total assemblies based on identity results from MashMap with minimum mapping region (i.e. “fragments”) length of 500bp. Each horizontal box is a scaffold of the reference on which the query scaffolds are mapped according to a given identity threshold. Mapped regions are ordered by base coordinate along the query scaffold on the x-axis, and the reference scaffolds on the y-axes. (A) CABOG assembly scaffold 1 mapped to the total polished Flye assembly, with corresponding Flye scaffold numbers reported on the right. (B) CABOG and Flye assemblies scaffold 1 mapped to the *S. chuatsi* and *C. auratus* chromosome-level assemblies.

### 3.3 Final assembly statistics

The Flye polished assembly provided the best results and thus was used in all subsequent analyses. This final genome assembly consisted of 567,902,715 bases in 1,214 scaffolds, with a scaffold N50 length of 3.37 Mb and a proportion of gaps of 0.001% (Table 2, Figure 6). Base composition was A: 28.10%, T: 28.15%, G: 21.87%, C: 21.88%, and overall standard deviation of GC content was 3.87%. The BUSCO completeness was very good overall, with more than 95% of the single-copy Actinopterygii orthologs retrieved in the final assembly (Table 2, Figure 6). The contiguity and completeness were high when compared to other Illumina + Nanopore hybrid assemblies (Table 3). The final assembly was named fNemMar1, in accordance with the Earth Biogenome Project sample naming scheme (https://gitlab.com/wtsi-grit/darwin-tree-of-life-sample-naming).

**Figure 6.**
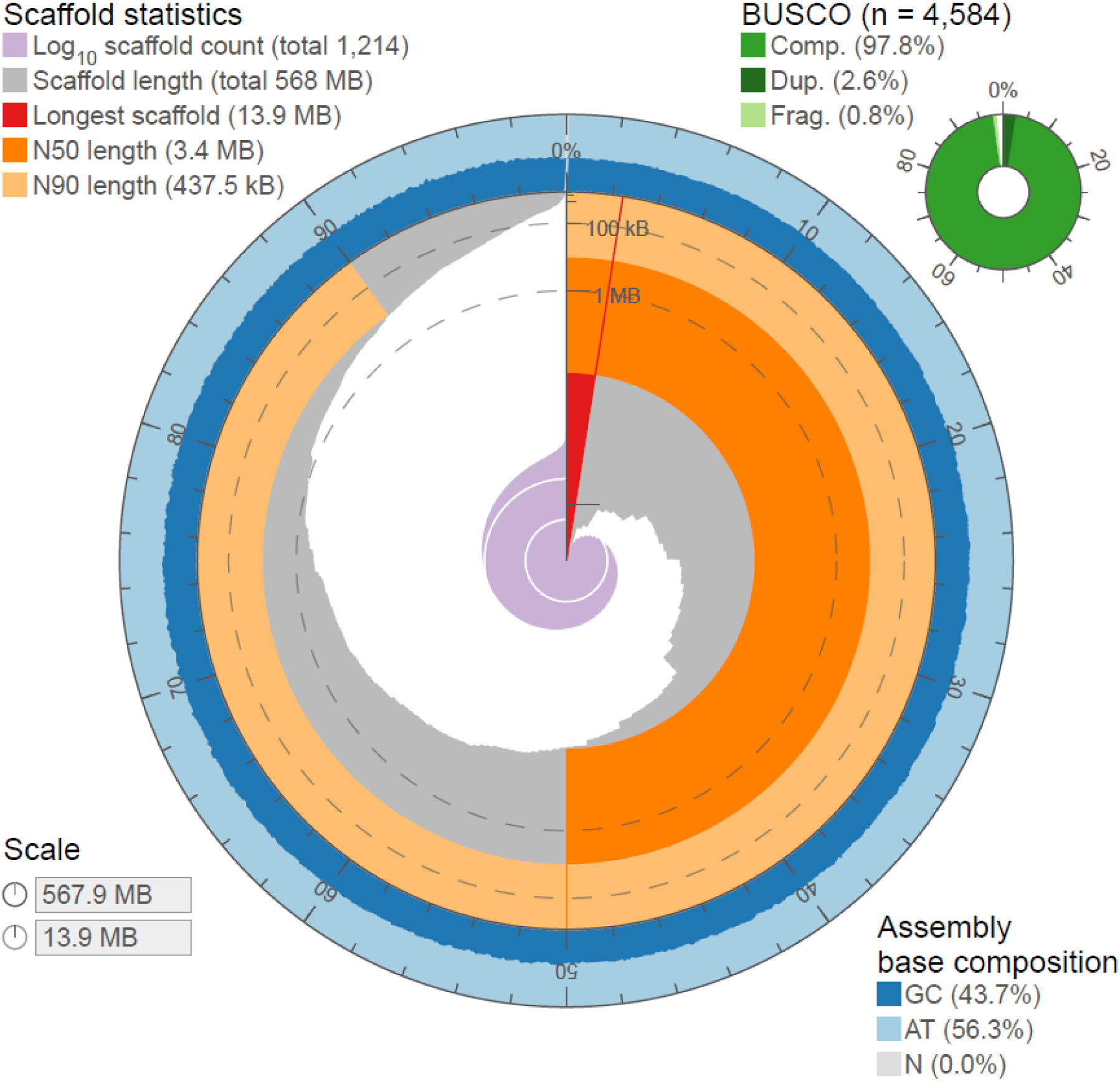
Visualization of contiguity and completeness of the final tarakihi assembly. The contiguity is visualized in a circle representing the full assembly length of c. 568 Mb. The longest scaffold was 13.9 Mb. There were very few scaffolds (c. 2%) shorter than 100 Kb in length and the GC content was uniform throughout. See Supplementary Figure 6 for a comparison with the three other assemblies that were not retained.

**Table 3.** Comparison of the contiguity and completeness of genomes that were assembled using a hybrid approach including only short Illumina reads and long Nanopore reads.

| Species | Genome (total scaffolds) length | Number of scaffolds | Scaffold N50 length | Complete BUSCOs | Protein-coding gene models | Functionally annotated genes |
| --- | --- | --- | --- | --- | --- | --- |
| Tarakihi | 568 Mb | 1,214 | 3.4 Mb | 97.80% | 20,327 | 19,823 |
| Murray cod | 633 Mb | 18,198 | 0.1 Mb | 94.20% | 26,539 | 25,607 |
| Clownfish | 881 Mb | 6,404 | 0.4 Mb | 96.30% | 27,420 | 26,211 |
| *Danionella translucida* | 735 Mb | 27,639 | 0.3 Mb | 91.50% | 24,097 | 21,491 |
| Snout otter clam | 544 Mb | 622 | 2.1 Mb | 95.80% | 26,380 | 23,701 |
| Indian blue peacock | 915 Mb | 15,025 | 0.2 Mb | not reported | 23,153 | 21,854 |

Note: All fish genome assemblies that corresponded to the criteria are reported (Murray cod (*Maccullochella peelii*): Austin et al. (2017), clownfish (*Amphiprion ocellaris*): Tan et al. (2018), *Danionella translucida*: Kadobianskyi et al. (2019)) and two selected additional species have been included for comparison with other groups of organisms (Mollusc, snout otter clam (*Lutraria rhynchaena*): Thai et al. (2019); bird, Indian blue peacock (*Pavo cristatus*): Dhar et al. (2019)).

### 3.4 Estimation of heterozygosity

Variant calling of Illumina reads against the polished assembly resulted in a total of 3,654,819 SNPs. By dividing this number by the size of the genome, this corresponded roughly to a heterozygosity level of 0.64%. This is lower than the level estimated by *k*-mer frequency (c. 1.00%). However, it is common for heterozygosity estimated by k-mer frequency to be lower compared to called SNPs, because the SNP calling approach is more conservative (Thai et al., 2019). Nevertheless, the heterozygosity estimated for TARdn1 is one of the highest reported for a fish species. To our knowledge, this is the highest heterozygosity estimated for a fish through *k*-mer analysis, with other reported values ranging from 0.1% (Tibetan loach *Triplophysa tibetana* and Murray cod *Maccullochella peelii*) to 0.9% (Java medaka *Oryzias javanicus*) (Austin et al., 2017; Ge et al., 2019; Gong et al., 2018; Lu et al., 2020; Nguinkal et al., 2019; Takehana et al., 2020; Vij et al., 2016; Yang et al., 2019; H. H. Zhang et al., 2020; Zheng et al., 2021). Even the heterozygosity estimated through SNPs (0.64%) is high compared to estimations from other fish using the same method (e.g. large yellow croaker: 0.36% (Wu et al., 2014), grass carp: 0.25% (Y. Wang et al., 2015)). This result is even more striking in that the variant analysis was very stringent in our case by retaining only high-quality bi-allelic SNPs. This reinforces the recent findings that *N. macropterus* is a species with a historically large population that displays a particularly high genetic diversity (Papa, Halliwell, et al., 2021).

### 3.5 Repetitive elements and genes annotation

REs represented 30.45% of the genome, or a total of 172,911,032 bp. Although the proportion of REs in fish genomes can vary greatly at scales from 10% to 60% (Yuan et al., 2018), the proportion of repeat elements in *N. macropterus* is on par with the proportion observed in other Centrarchiformes (Largemouth bass (*Micropterus salmoides*): 33.79%, Big-eyed mandarin fish (*Siniperca knerii*): 26.55%) (Lu et al., 2020; Sun et al., 2021) and for Perciformes in general (Yuan et al., 2018). Of the REs known in the databases, interspersed repeats accounted for 27.62% of the genome, including 10.87% of DNA transposons and 6.17% of retro-elements (LINEs, LTR, SINEs, and PLE in that order). The rest of the repeat elements consisted of simple sequence repeats (Supplementary Table 1).

After filtering for length, the final predicted gene set included 20,169 protein-coding genes with a mean length of 13,832 bp, among which 95.5% had an AED < 0.5. The mean exon length was 229 bp, and the mean intron length in CDS was 1,184 bp. More than 98% of the genes were functionally annotated by at least one of the two methods used (blastp 98.2%, InterProScan 82.8%).

### 3.6 Iso-Seq analysis

Of the 93,949 full-length polished, non-redundant Iso-Seq transcripts, 2,347 were classified as REs and were filtered out from downstream analyses. For each of these RE transcripts, the main RE elements included DNA elements (801), LINEs (639), LTRs (464), SINEs (94), rRNAs (47), low complexity / simple repeats (33), rolling circles (26), satellites (16), and retroposons (2), as well as one LINE/LTR hybrid, and 224 unknown RE.

The final non-RE Iso-Seq dataset included 91,313 unique transcripts from 15,515 genes. The mean transcript per gene ratio was 5.89, with a median of 3 and a maximum of 211 (Figure 7A). This is higher than the values recently reported for humans (3.62) and two species of bats (1.92 and 1.49), but lower than pharaoh ants (9) (Q. Gao et al., 2020; Wen et al., 2018). Less than 5% of genes had more than 20 different transcripts. The predicted proteins of both genes that produced the most transcripts (respectively 211 and 164 transcripts) were collagen alpha chains isoforms (XP_006787735.1: collagen alpha-2(I) chain-like isoform X2, XP_020490299.1: collagen alpha-1(V) chain-like isoform X1), implicated in the structural integrity of the cellular matrix (GO:0005201).

**Figure 7.**
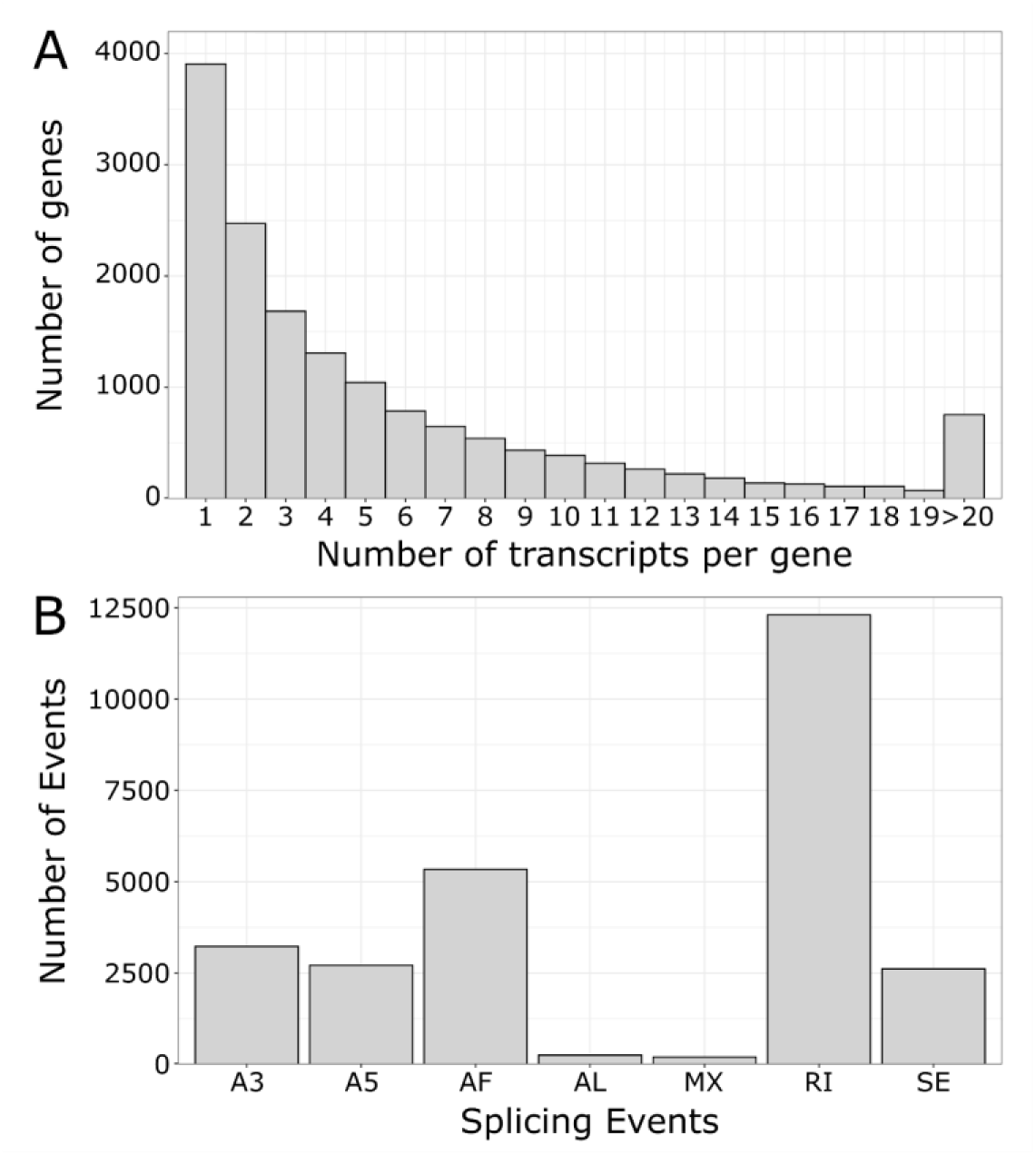
Alternative transcripts metrics in the tarakihi transcriptome (A) Number of unique alternative transcripts per gene. (B) Classification and frequency of alternative splicing events. A5/A3: Alternative 5’/3’ Splice Sites. AF/AL: Alternative First/Last Exons. MX: Mutually Exclusive Exons. RI: Retained Intron. SE: Skipping Exon.

A total of 26,644 AS events were detected in the tarakihi transcriptome (Figure 7B). The most frequent AS event was the retention of intron (46%), while “alternative last exons” and “mutually exclusive exons” were the rarest (less than 1% each). Some examples of these AS events were visualized in the tarakihi genome (Figure 8).

**Figure 8.**
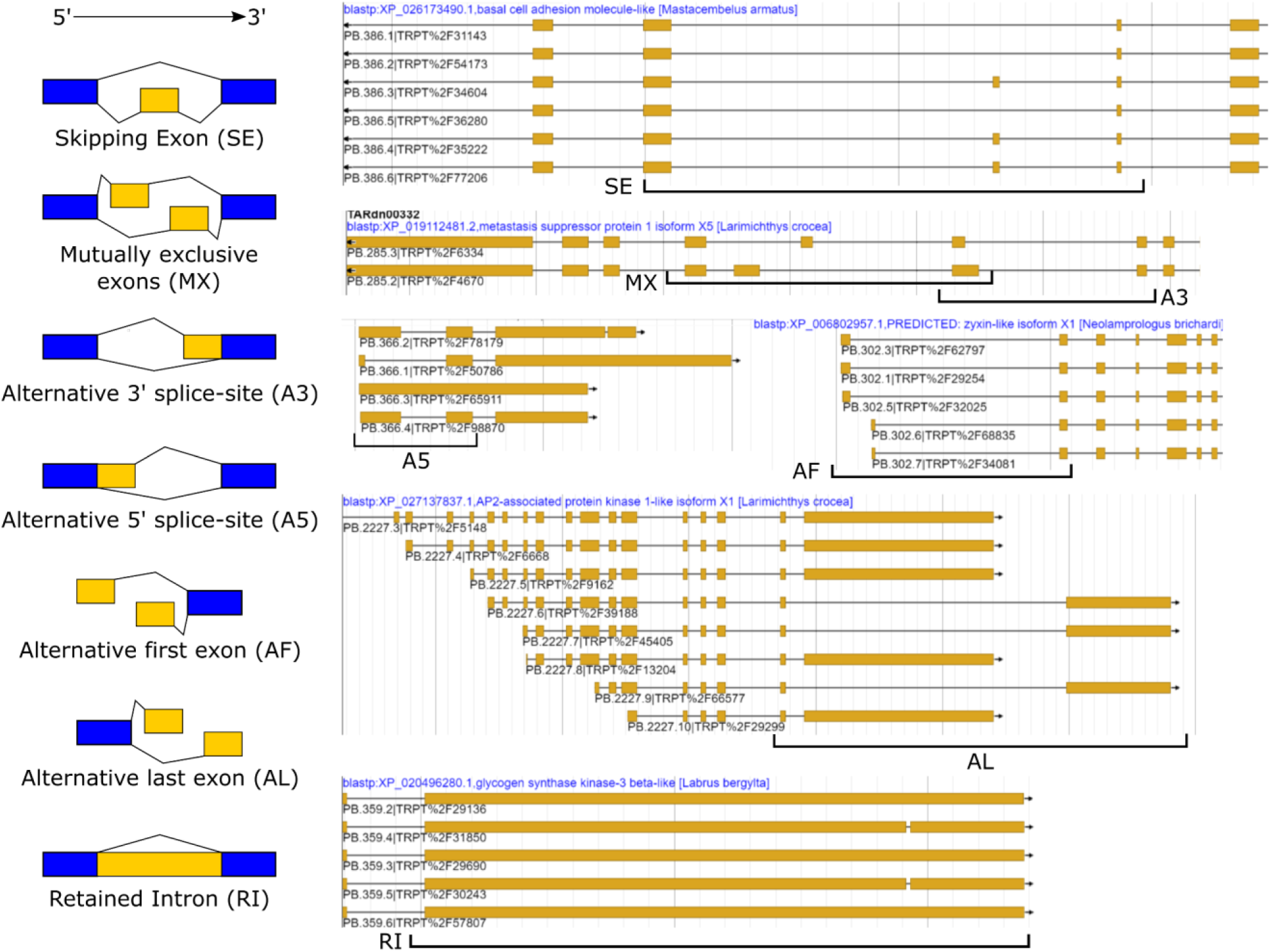
The seven types of alternative splicing events classified in the tarakihi transcriptome, with examples of each event class as visually shown in the annotation of the genome.

Comparison of the frequency of AS events in the tarakihi with other species showed that the trends are globally similar across organisms (Figure 9). Most organisms show relatively high occurrences of RI, A3, A5, AF, and to a lesser degree SE, compared to AL and MX. The figure also shows that tarakihi, goldfish, and cave nectar bat may have a better representation of the AS events proportions due to a much deeper coverage compared to the Wuchang bream, zebrafish, and whiteleg shrimp (although values for MX and SE were not reported for the goldfish study). While it is the most common AS event in both tarakihi and goldfish, the proportion of RI events is much higher in tarakihi compared to the proportion of other events. While intron retention was thought until recently to be the least prevalent AS form in animals, it is now clear that this is not the case (as shown in the studies in Figure 9 but also e.g. Q. Gao et al. (2020); X. Wang et al. (2019)). RI events are widely used across organisms to tune down the levels of transcription of some genes in cells and tissues depending on their function (Braunschweig et al., 2014).

**Figure 9.**
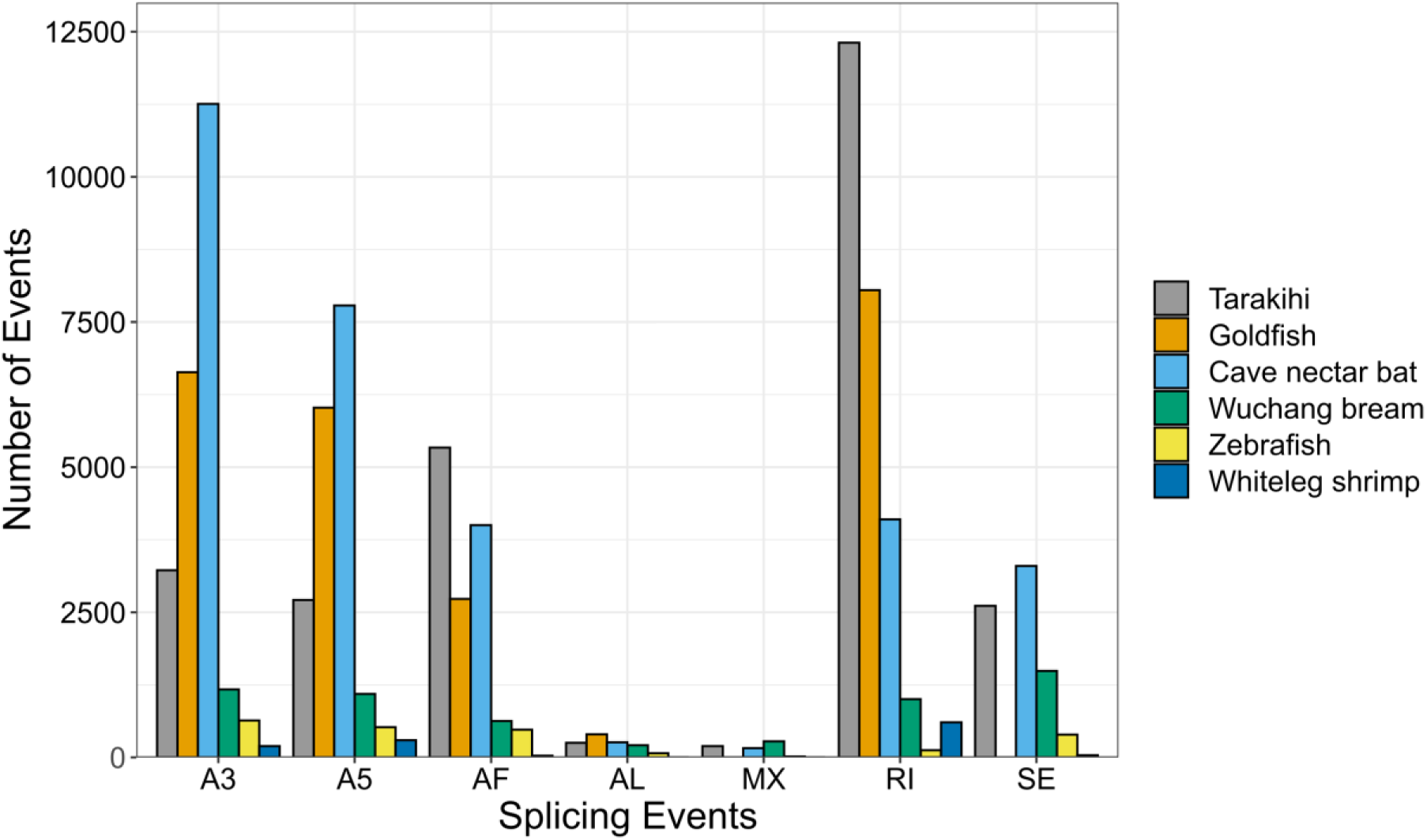
Comparison of alternative splicing event counts between tarakihi and five other animal species from other Iso-Seq AS studies. MX and SE events were not reported in the goldfish study.

### 3.7 Genome size

The size of the tarakihi genome was consistent with values for fish genomes that have been reported so far. A recent review of publicly available fish genome assemblies (comprising 244 species) showed that the average genome length of fish is 872.64 Mb but varies between c. 300 Mb to c. 4.5 Gb (Fan et al., 2020). The genome size of *N. macropterus* (568 Mb) is several hundred Mb shorter than the two other published Centrarchiforme genomes, the largemouth bass *Micropterus salmoides* (964 Mb) and the big-eye mandarin fish *Siniperca knerii* (732.1 Mb) (Lu et al., 2020; Sun et al., 2021). However, *N*. *macropterus* is still evolutionarily far apart from these two species. The largemouth bass and the big-eye mandarin fish both belong to the Centrarchoidei sub-order, which is thought to have split from Cirrhitioidei at least 70 million years ago (Sanciangco et al., 2016).

## 4. Conclusion

The advances in DNA sequencing technologies have made it clear how valuable reference genome assemblies are for the study of biology and conservation, resulting in a global effort to assemble the genomes of as many organisms as possible (Fan et al., 2020; Koepfli et al., 2015; Worley et al., 2017). Here we present the first genome assembly of the tarakihi, a valuable commercial fisheries species, and the first representative out of the c. 60 species of the Cirrhitioidei suborder to have a whole genome sequenced. While performing a hybrid assembly of Illumina and Nanopore reads with the latest tools led to a highly contiguous assembly with high gene completeness, this could be still improved in the future by adding Hi-C data to scaffold it to a chromosome-level assembly (Whibley et al., 2021). Moreover, while PacBio HiFi data was a very new and still relatively expensive technology at the time of data collection, it will probably replace the short and long reads hybrid assembly method as the optimal genome assembly strategy by offering the best of both worlds (long reads and high quality) and allowing phasing of genomes. However, the present genome and its accompanying highly accurate transcriptome will still be a valuable resource for future studies, including, but not restricted to comparative genomics, population structure analyses, and the study of adaptive selection.

## 5. Acknowledgments

The authors are thankful to Igor Ruza, David Ashton, and Matthew Wylie (Plant and Food Research, Nelson) for assisting in the sampling of the captive-bred specimen and to Nick Johnston for capturing and providing the wild specimen. They are also grateful for advice and assistance from the Removing Fisheries Juvenile Habitat Bottleneck Technical Advisory group, and in particular, they wish to acknowledge the support and consultation from Laws Lawson (Te Ohu Kaimoana), Jeremy Helson (Fisheries Inshore New Zealand), and Carol Scott (Southern Inshore Fisheries Management Company Limited). They thank Tom Oosting and Holly Jackson (Victoria University of Wellington) for proofreading the manuscript.

## 6. CRediT authorship contribution statement

**Yvan Papa:** Conceptualisation, Methodology, Software, Validation, Formal analysis, Investigation, Resources, Data Curation, Writing - Original Draft, Writing - Review & Editing, Visualisation. **Maren Wellenreuther:** Resources, Writing - Review & Editing, Supervision, Funding acquisition. **Mark A. Morrison:** Writing - Review & Editing, Supervision, Funding acquisition. **Peter A. Ritchie:** Conceptualisation, Resources, Writing - Review & Editing, Supervision, Project administration, Funding acquisition.

## 9. Data availability statement

All genomic sequences and associated metadata are deposited on the Genomics Aotearoa repository (https://repo.data.nesi.org.nz/) under project name “tarakihi genomics”. All scripts used in the analyses are openly available on GitHub at https://github.com/yvanpapa/tarakihi_genome_assembly.

## 11. Supplementary Material

**Supplementary Figure 1.**
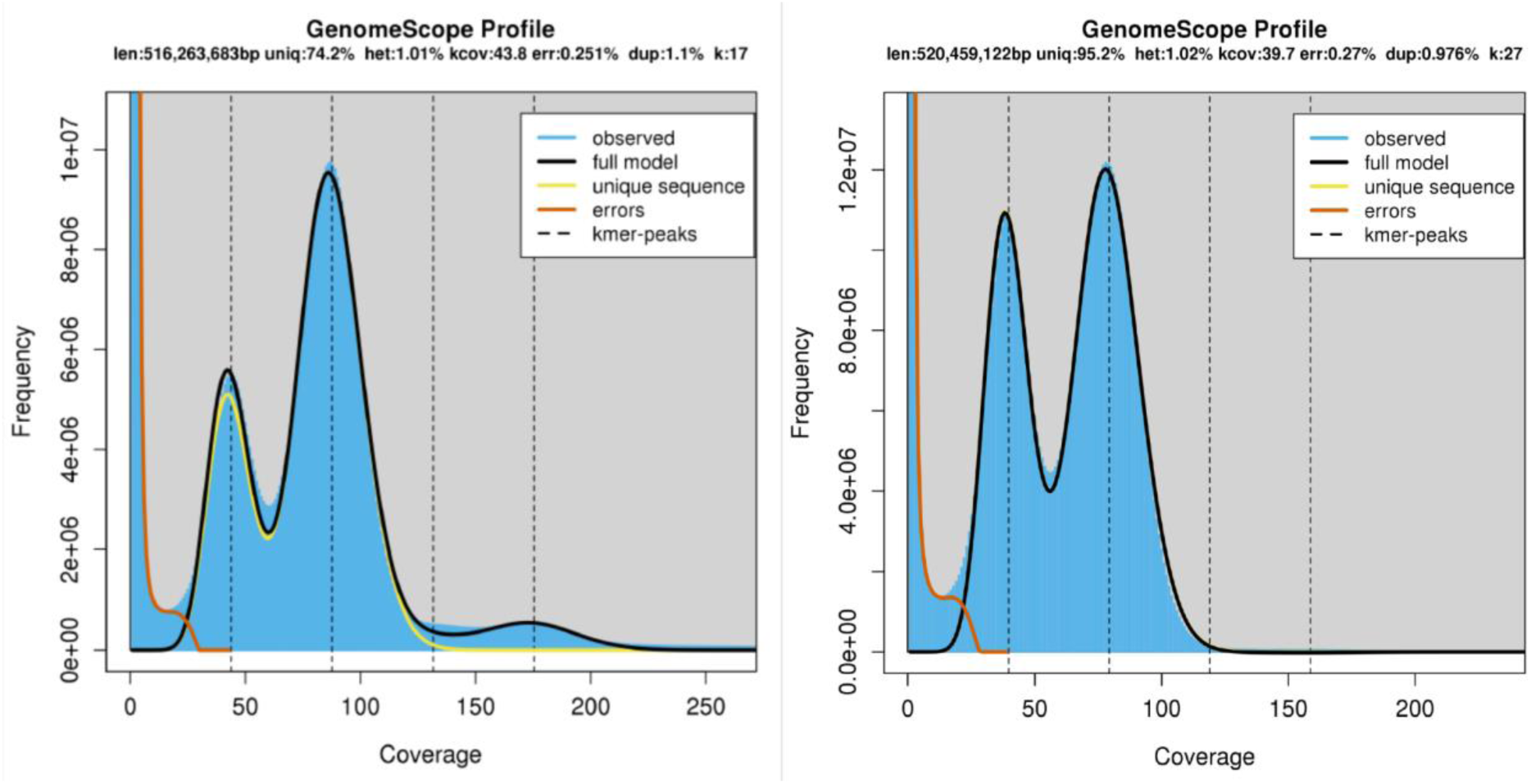
Histograms of 17- and 27-mer frequency in clean Illumina reads.

**Supplementary Figure 2.**
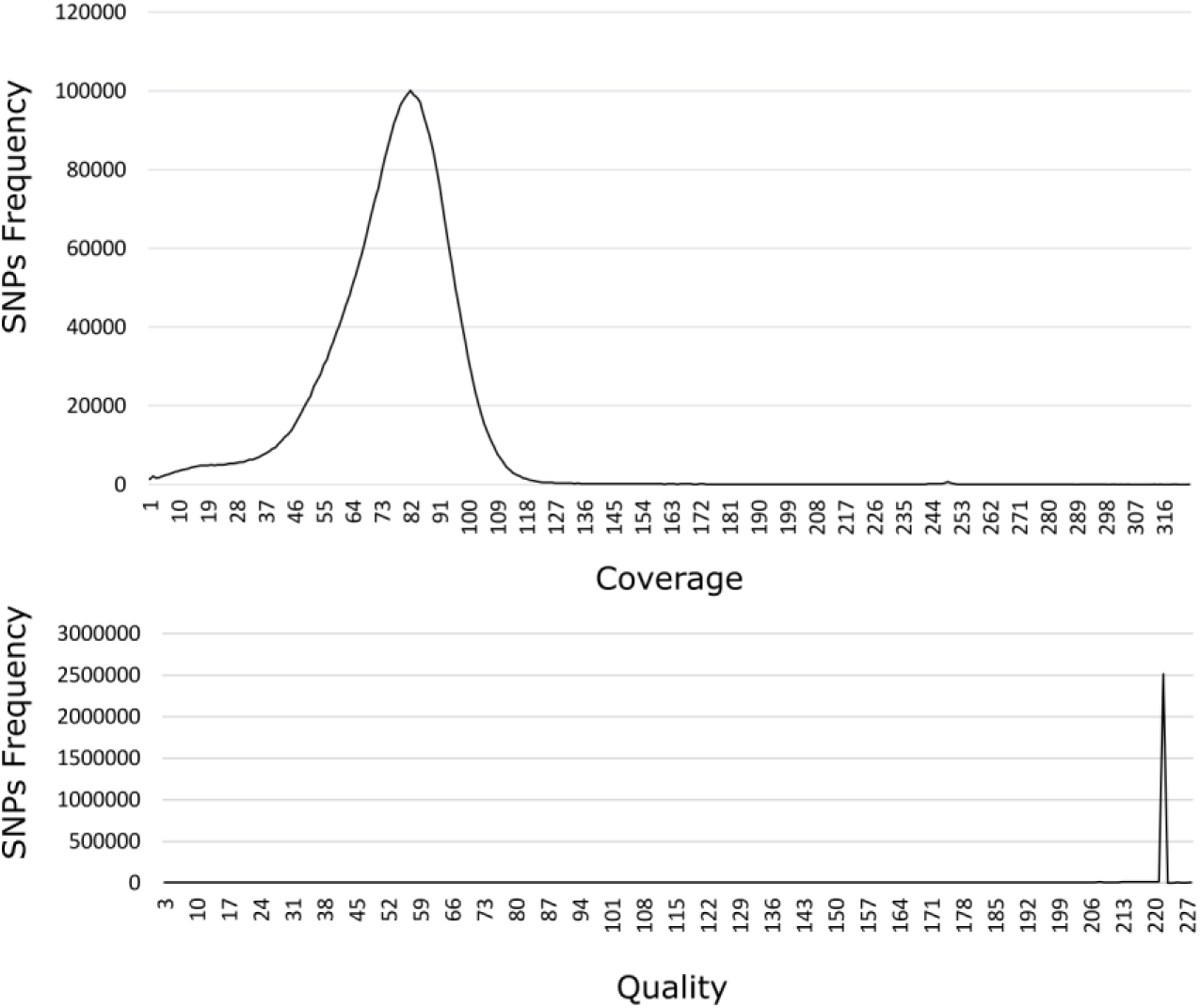
Distribution of coverage (top) and quality (bottom) of SNPs called from Illumina reads back to the assembly. SNPs were filtered for a minimum genotype depth of 20 according to the increase in steepness starting approximately at this point. Quality was always high, so the default site quality value was used.

**Supplementary Figure 3.**
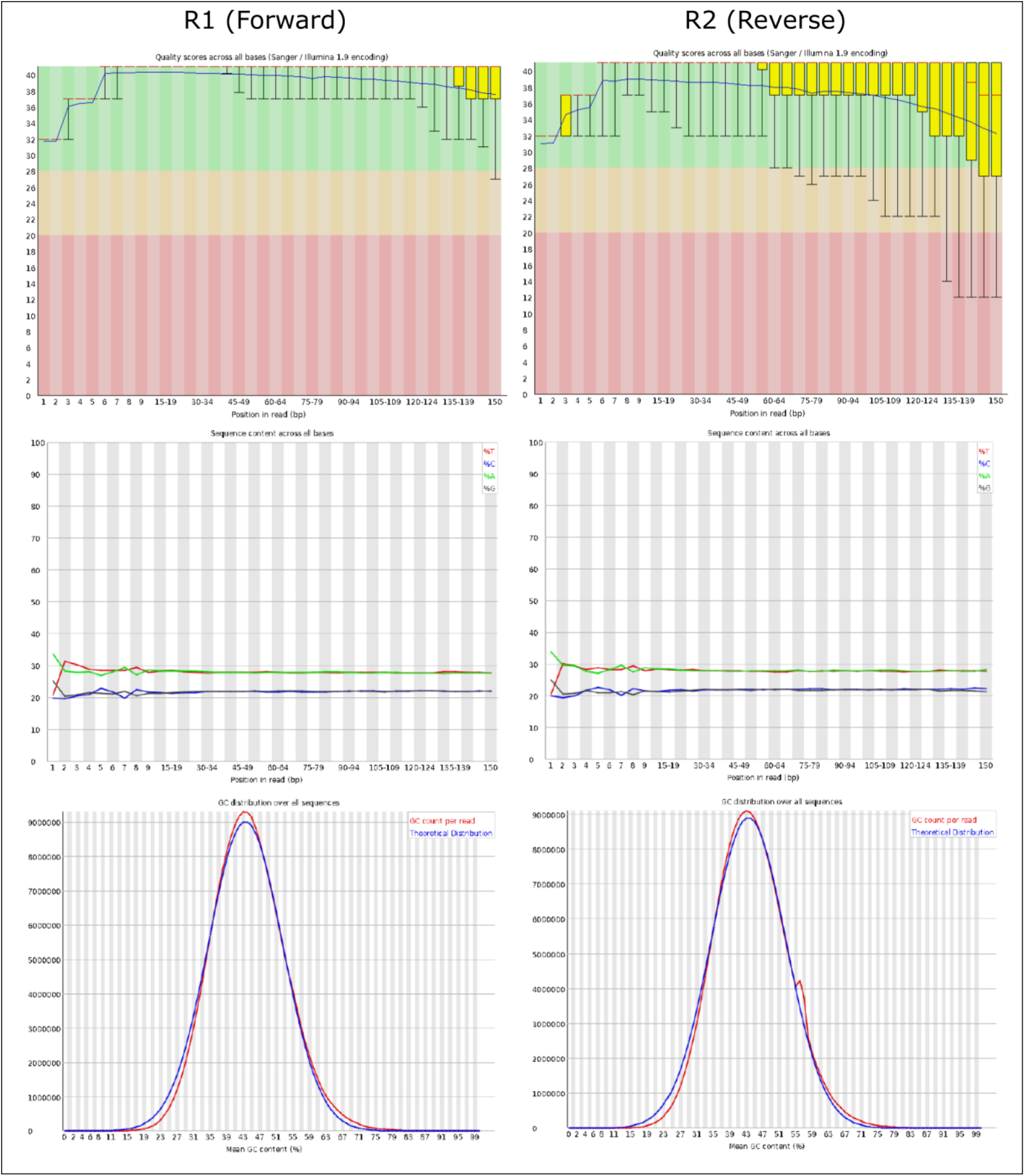
Some of FastQC quality metrics results, for forward (R1) and reverse (R2) reads. Top: Per base sequence quality. Middle: Per base sequence base content. Bottom: GC distribution over all sequences. See main text for the explanation on the slight bias in bases content for the first few bases in all reads. GC content of reverse reads detected a few over-represented sequences, which were most probably harmless sequencing artifacts that should be discarded during the quality control step of the MaSuRCA assembly.

**Supplementary Figure 4.**
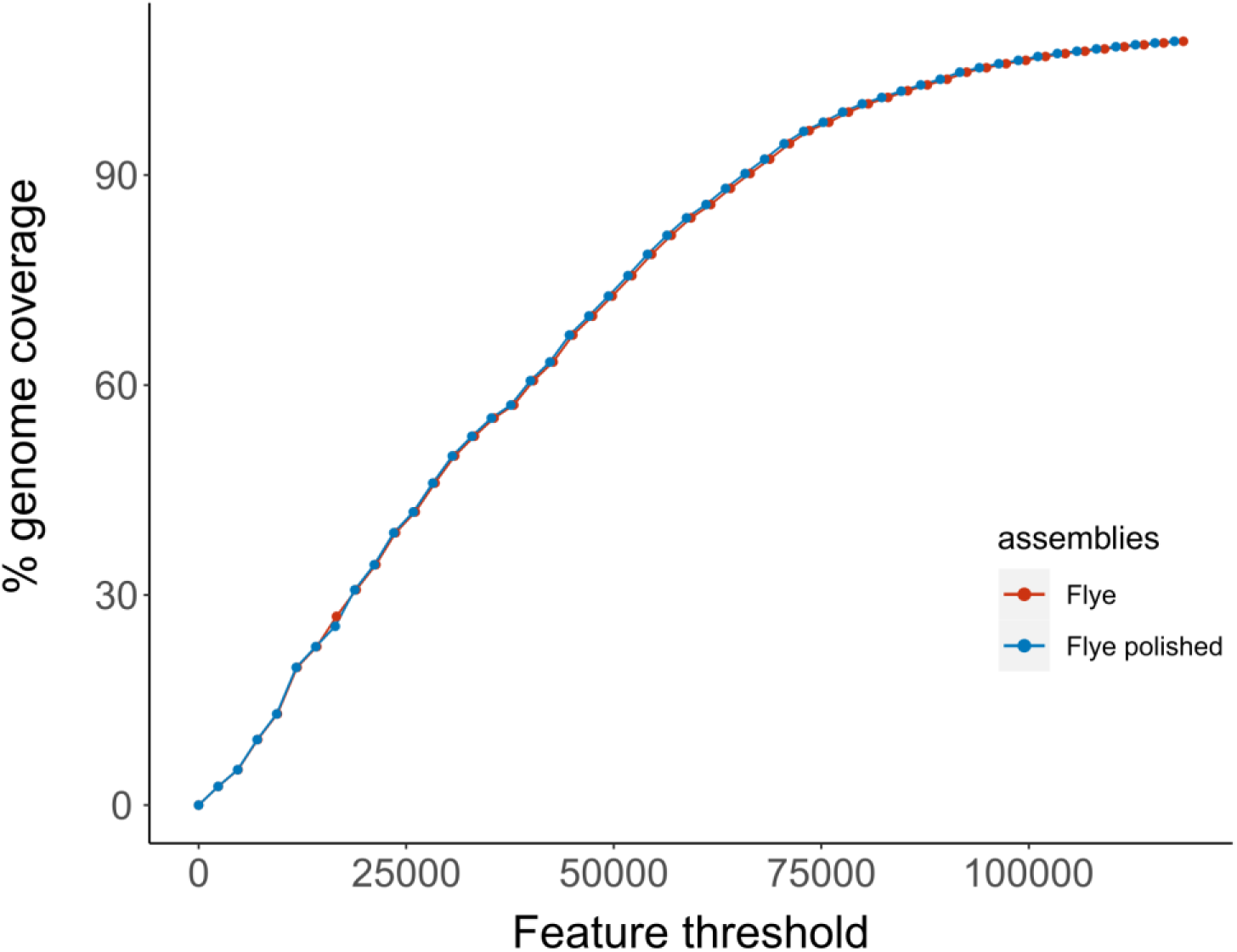
FRC curves as shown in Figure 3, but with only the Flye and Flye polished assemblies projected for better visualization. For the same cumulative genome size, the Flye unpolished assembly always accumulates slightly more potential errors (i.e. features).

**Supplementary Figure 5.**
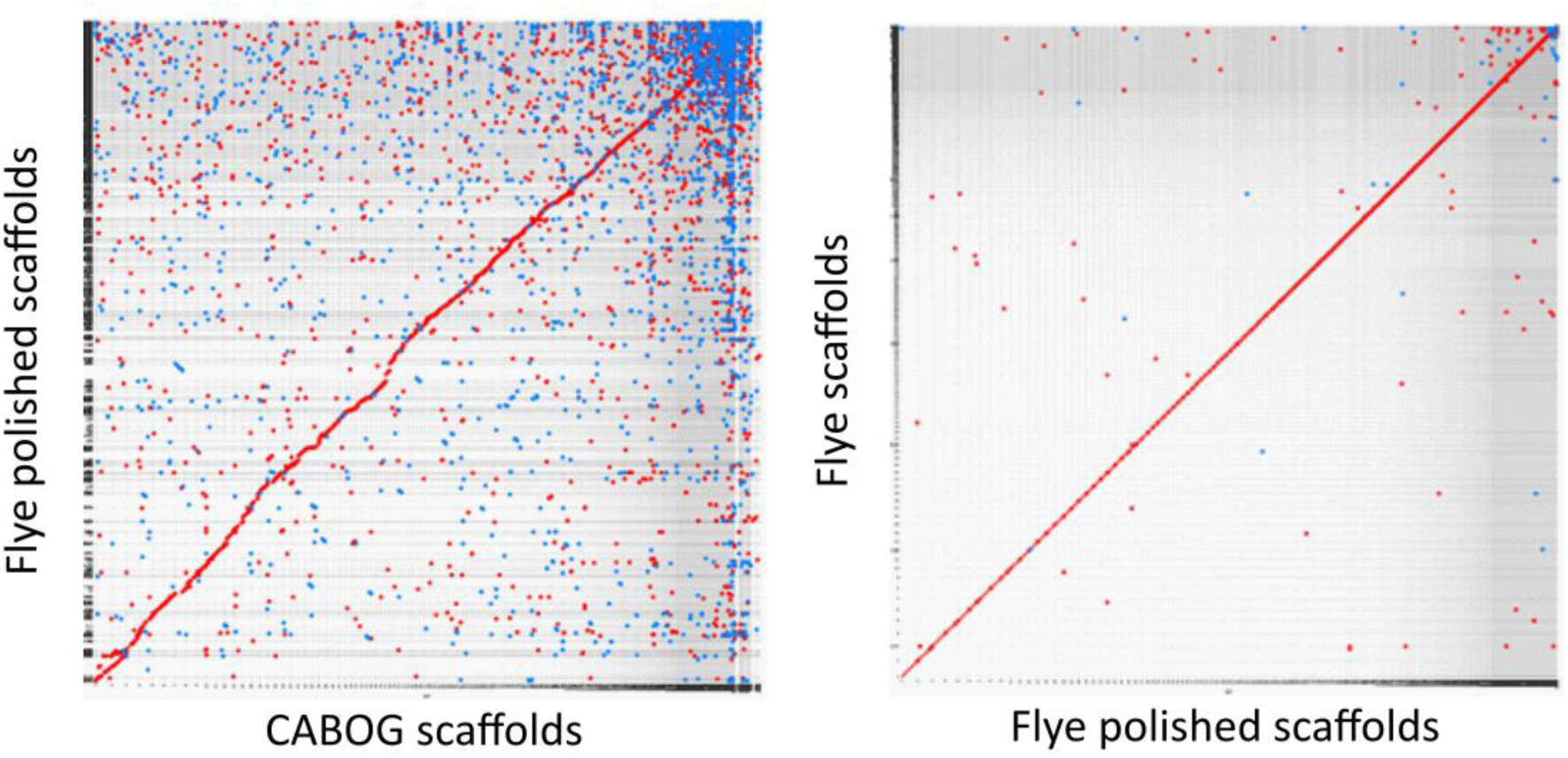
Plots of pairwise alignment scores between scaffolds, obtained with MashMap. Each dot represents a match between the query and the reference sequence. Colors correspond to the strand direction (red for positive, blue for negative).

**Supplementary Figure 6.**
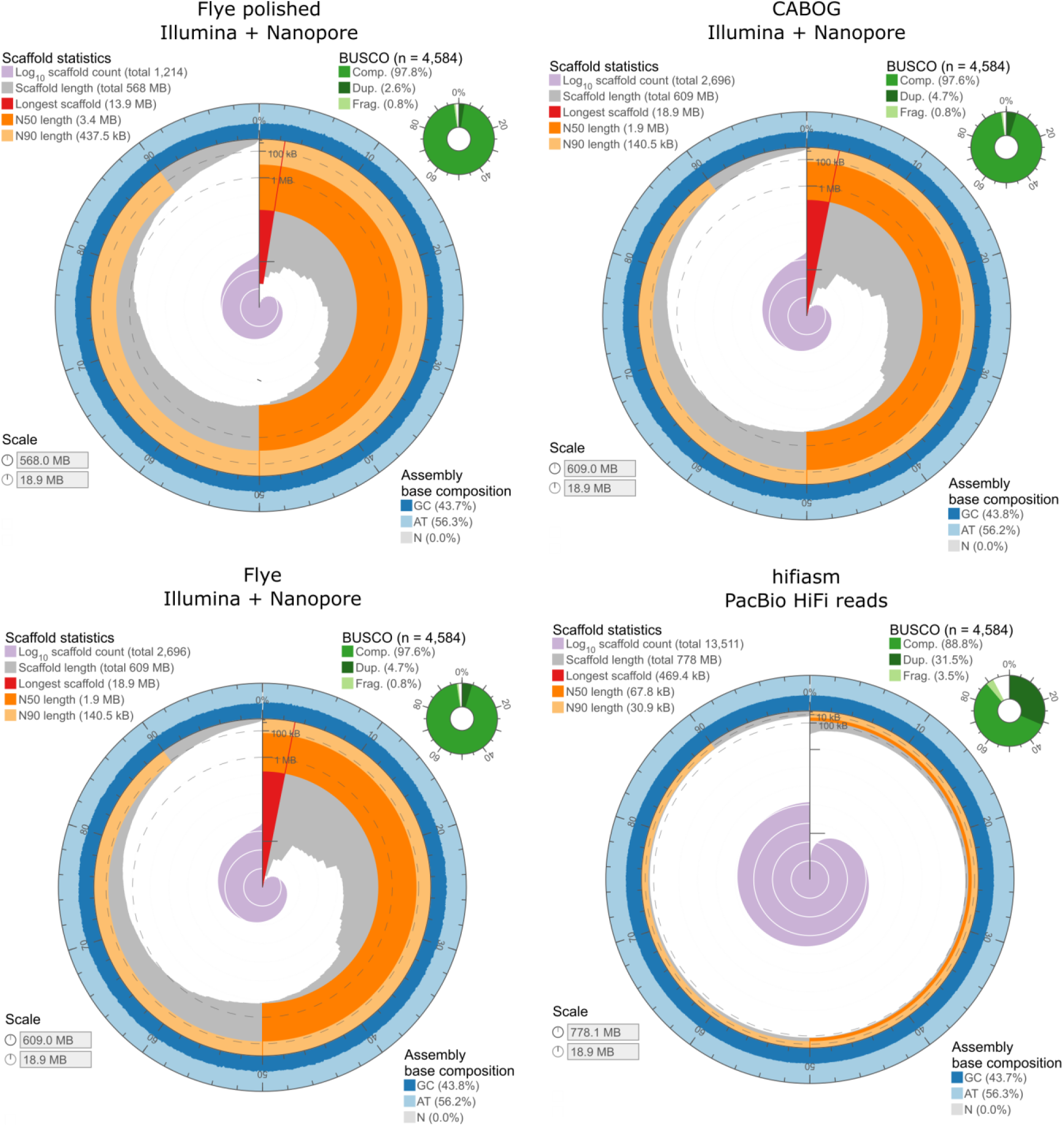
Visualization of contiguity and completeness of the four assemblies produced.

**Supplementary Table 1.**
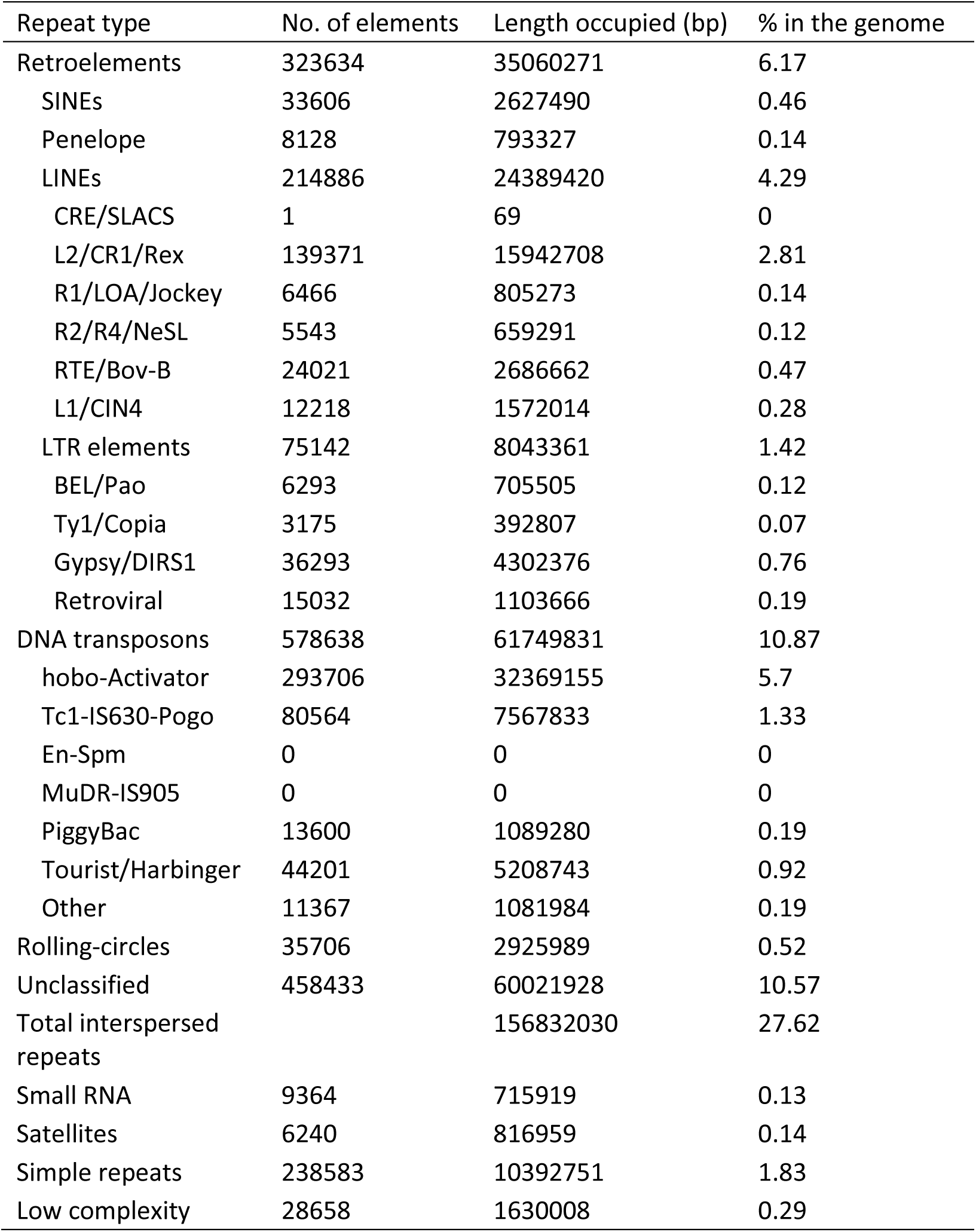
Main classes and proportions of repeat elements detected in the tarakihi genome.

| Repeat type | No. of elements | Length occupied (bp) | % in the genome |
| --- | --- | --- | --- |
| Retroelements | 323634 | 35060271 | 6.17 |
| SINEs | 33606 | 2627490 | 0.46 |
| Penelope | 8128 | 793327 | 0.14 |
| LINEs | 214886 | 24389420 | 4.29 |
| CRE/SLACS | 1 | 69 | 0 |
| L2/CR1/Rex | 139371 | 15942708 | 2.81 |
| R1/LOA/Jockey | 6466 | 805273 | 0.14 |
| R2/R4/NeSL | 5543 | 659291 | 0.12 |
| RTE/Bov-B | 24021 | 2686662 | 0.47 |
| L1/CIN4 | 12218 | 1572014 | 0.28 |
| LTR elements | 75142 | 8043361 | 1.42 |
| BEL/Pao | 6293 | 705505 | 0.12 |
| Ty1/Copia | 3175 | 392807 | 0.07 |
| Gypsy/DIRS1 | 36293 | 4302376 | 0.76 |
| Retroviral | 15032 | 1103666 | 0.19 |
| DNA transposons | 578638 | 61749831 | 10.87 |
| hobo-Activator | 293706 | 32369155 | 5.7 |
| Tc1-IS630-Pogo | 80564 | 7567833 | 1.33 |
| En-Spm | 0 | 0 | 0 |
| MuDR-IS905 | 0 | 0 | 0 |
| PiggyBac | 13600 | 1089280 | 0.19 |
| Tourist/Harbinger | 44201 | 5208743 | 0.92 |
| Other | 11367 | 1081984 | 0.19 |
| Rolling-circles | 35706 | 2925989 | 0.52 |
| Unclassified | 458433 | 60021928 | 10.57 |
| Total interspersed repeats |  | 156832030 | 27.62 |
| Small RNA | 9364 | 715919 | 0.13 |
| Satellites | 6240 | 816959 | 0.14 |
| Simple repeats | 238583 | 10392751 | 1.83 |
| Low complexity | 28658 | 1630008 | 0.29 |

**Supplementary Table 2.** Quality control statistics of the gene models obtained after different rounds of MAKER.

|  | Round 1 | Round 2 | Round 3 |
| --- | --- | --- | --- |
| Number of gene models | 9,008 | 20,327 | 19,930 |
| Average gene length | 11,455 | 13,741 | 14,057 |
| AED ≤ 0.5 | 100% | 95.50% | 94.00% |
| Complete BUSCO transcripts | 58.70% | 76.00% | 74.90% |

